# Klf5 establishes bi-potential cell fate by dual regulation of ICM and TE specification genes

**DOI:** 10.1101/2021.06.02.446799

**Authors:** Martin Kinisu, Yong Jin Choi, Claudia Cattoglio, Ke Liu, Hector Roux de Bezieux, Raeline Valbuena, Nicole Pum, Sandrine Dudoit, Haiyan Huang, Zhenyu Xuan, Sang Yong Kim, Lin He

**Affiliations:** Division of Cellular and Developmental Biology, MCB department, University of California at Berkeley, Berkeley, CA 94705, USA; Howard Hughes Medical Institute, University of California, Berkeley, Berkeley, CA 94720, USA; Department of Statistics, University of California at Berkeley, Berkeley, CA 94720, USA; Division of Biostatistics, School of Public Health, University of California at Berkeley, Berkeley, CA 94720, USA; Department of Molecular and Cell Biology, University of Texas at Dallas, 800 West Campbell Road, Richardson, Texas 75080, USA; Department of Pathology, NYU Grossman School of Medicine, 540 First Ave, New York, NY 10016, USA

## Abstract

Early blastomeres of mouse preimplantation embryos exhibit bi-potential cell fate, capable of generating both embryonic and extra-embryonic lineages in blastocysts. Here, we identified three major 2 cell (2C) specific endogenous retroviruses (ERVs) as the molecular hallmark of the bi-potential plasticity. Using the LTRs of all three 2C-ERVs, we identified Klf5 as their major upstream regulator. Klf5 is essential for bi-potential cell fate: a single *Klf5-overexpressing* ESC generated terminally differentiated embryonic and extra-embryonic lineages in chimeric embryos, and Klf5 directly induces both ICM and TE specification genes. Intriguingly, *Klf5* and *Klf4* act redundantly during ICM specification, whereas *Klf5* deficiency alone impairs TE specification. *Klf5* is regulated by multiple 2C-specific transcription factors, particularly Dux, and the Dux/Klf5 axis is evolutionarily conserved. Altogether, the 2C-specific transcription program converges on Klf5 to establish bi-potential cell fate, enabling a cell state with dual activation of ICM and TE genes.

## Introduction

Mammalian preimplantation development is initiated by maternally inherited factors and zygotic genes transcribed during zygotic genome activation (ZGA)(Deng et al., 2014). Mouse zygotes and 2C blastomeres are totipotent, capable of generating all cell types requisite for a fertile adult organism (Casser et al., 2017). Totipotency is gradually restricted in subsequent developmental stages (4C to 8C stage), (Wu et al., 2017); yet cleavage stage blastomeres retain bi-potential cell fate, generating the inner cell mass (ICM) that largely forms the embryo proper, and trophectoderm (TE) that gives rise to extra-embryonic placental tissues(Korotkevich et al., 2017), (Fujimori et al., 2003; Tabansky et al., 2013), (Wigger et al., 2017).

A prominent molecular hallmark of bi-potential cell fate is a strong yet transient induction of MERVL endogenous retroviruses (ERVs)(Choi et al., 2017; Macfarlan et al., 2011, 2012), (Ishiuchi et al., 2015a). MERVL transcripts are among the most highly expressed transcripts in the transcriptomes of 2C-4C blastomeres (Franke et al., 2017); they quickly decrease in level as the developmental plasticity of cleavage stage blastomeres narrows during development. In pluripotent mouse embryonic stem cells (ESCs), which generate all embryonic cell types but rarely extraembryonic lineages (Beddington and Robertson, 1989), MERVL induction in rare cell populations often correlates with an expanded cell fate plasticity, enabling differentiation towards both embryonic and extra-embryonic lineages (Choi et al., 2017; Ishiuchi et al., 2015a; Macfarlan et al., 2011, 2012; Schoorlemmer et al., 2014; Zhao et al., 2018)(Hu et al., 2020; Yan et al., 2019). However, such MERVL^+^ ESCs are not equivalent to 2C blastomeres (Choi et al., 2017), possessing neither totipotent cell fate potential, nor a 2C-transcriptome. Rather, the MERVL^+^ ESCs, designated as bi-potential ESCs in our study, exhibit both embryonic and extraembryonic potency, induce MERVL at a modest level, and functionally resemble bi-potential blastomeres, which are more restricted in developmental potentials than 2C blastomeres.

Since MERVL is a molecular hallmark of 2C blastomeres, transcription factors that directly regulate MERVL and transiently peak at ZGA have been speculated to establish a transcriptome landscape of bi-potential cell fate (Alda-Catalinas et al., 2020; Eckersley-Maslin et al., 2019; Hendrickson et al., 2017; Iaco et al., 2017; Yan et al., 2019). The double homeodomain transcription factor Dux is one such candidate, induced at the onset of ZGA to directly promote MERVL expression in 2C blastomeres (Eckersley-Maslin et al., 2019; Hendrickson et al., 2017; Iaco et al., 2017). Zygotically expressed Dux, as well as its maternally inherited upstream regulators, Dppa2, Dppa4, Nelfa and Smarca5, have all been speculated to act at the top of the transcriptional hierarchy that governs the onset of ZGA, induces MERVL and regulates 2C-specific cell fate potency (Alda-Catalinas et al., 2020; Eckersley-Maslin et al., 2019; Hu et al., 2020). Yet deficiency of *Dux*, *Smarca5* or *Nelfa* alone, or *Dppa2 and Dppa4* in combination, fails to impair ICM or TE specification in mice, suggesting that these factors are not essential to establish/maintain bi-potential cell fate (Chen and Zhang, 2019; Nakamura et al., 2011), ((IMPC), 2014), (Stopka and Skoultchi, 2003). Hence, using MERVL as the sole molecular marker for bi-potential cell fate may not be sufficient to identify the key regulator(s) for bi-potential developmental cell fate.

Using three 2C specific ERVs as the hallmark of bi-potential cell fate, we identified Klf5 as an essential regulator that confers a developmental potency to both embryonic and extraembryonic lineages. While previous studies have characterized the preimplantation defects of *Klf5* knockout embryos(Azami et al., 2017; Ema et al., 2008; Lin et al., 2010), the functional importance of Klf5 in bi-potential cell fate and the functional interaction between Klf5 and other Klf transcription factors remain largely unknown. Here, our studies showed that *Klf5* overexpression in a single, pluripotent ESC confers a bi-potential cell fate in chimeric embryos. Our mouse genetics studies demonstrated the essential role of *Klf5* in enabling TE specification, and the redundant role of *Klf5* and *Klf4* in conferring potency for the ICM cell fate. Since Klf5 directly induces both ICM and TE specification genes(Lin et al., 2010), our data suggest that the molecular nature of a bi-potential developmental potency is a cell state that co-expresses ICM and TE genes. While

## Results

### Identification of 2C-specific ERV families

We set out to comprehensively identify 2C-specific retrotransposon markers in search of a master transcriptional regulator of bi-potential cell fate (Fig. 1A). Using published single cell RNA-seq data on preimplantation embryos, we observed a dynamic and tightly regulated expression pattern of retrotransposons (Fig. 1B, Sup Fig. S1A). While MERVL induction is a prominent molecular hallmark of 2C blastomeres, two additional ERVs, ORR1A0 and ORR1A1, also constitute the major retrotransposon families with a 2C-specific expression pattern (Fig. 1B, 1C, Sup Fig. S1A). At the peak of their expression, MERVL, ORR1A0, and ORR1A1 each account for 5.3%, 1.5% and 1.1% of all mapped reads in 2C blastomeres, respectively (Sup Fig. S1A). Subsequently, all three ERV families rapidly declined by the 8C stage, and were completely silenced in blastocysts (Fig. 1B, 1C, Sup Fig. S1A). Their coordinated induction was confirmed in a subset of bi-potential, MERVL+ ESC lines (Fig. 1D). Specifically, *Lsd1* deletion, and knockdown of Caf-1 subunits *P60* and *P150* in ESCs all yield coordinated induction of these three 2C-specific ERVs (Fig. 1D). Since MERVL, ORR1A0, and ORR1A1 collectively marked the transcriptional state of 2C blastomeres and bi-potential ESCs, we hypothesized that a transcription factor(s) capable of inducing all three 2C-specific ERVs could functionally confer 2C-like, bi-potential cell fate.

**Figure. 1.**
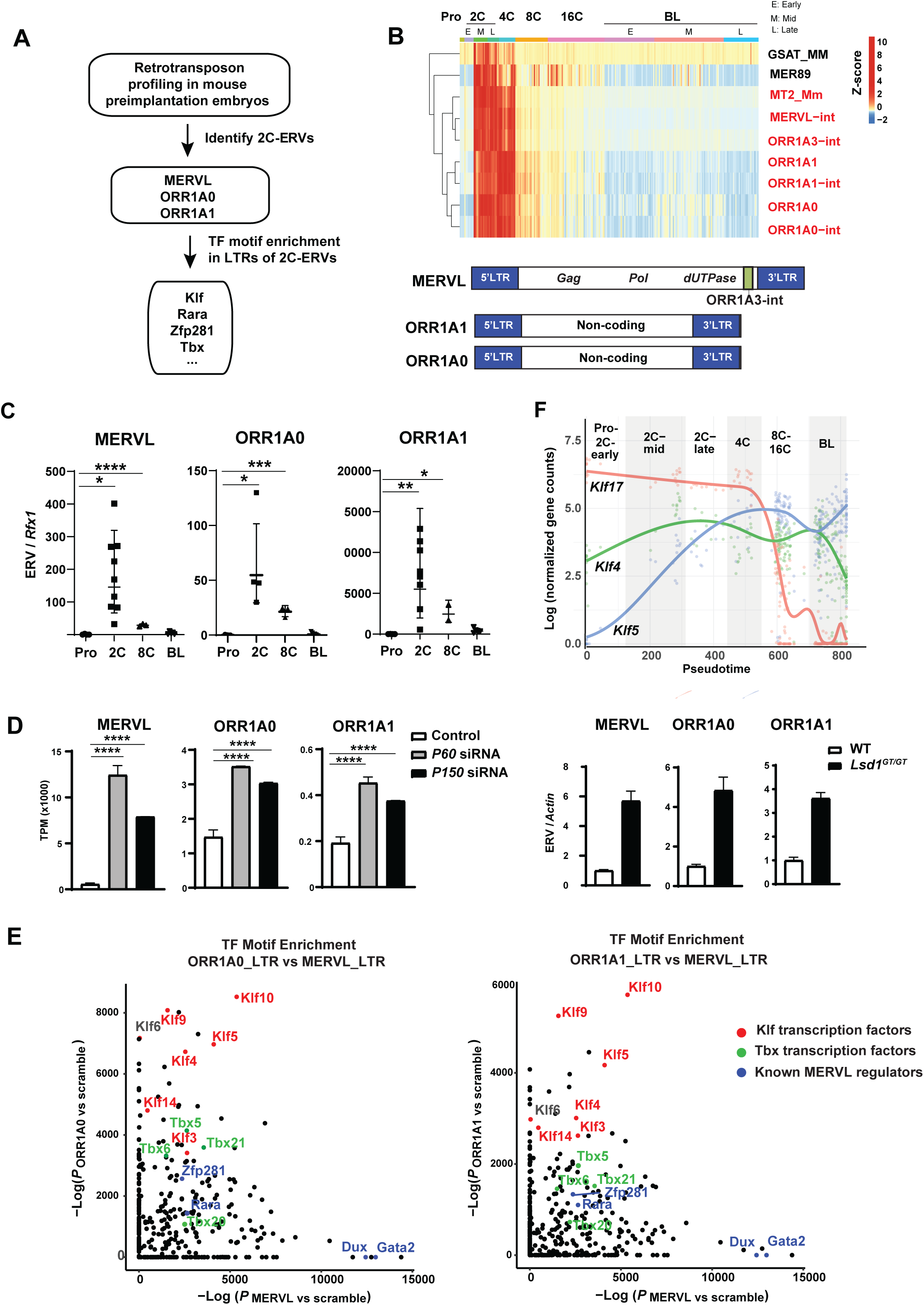
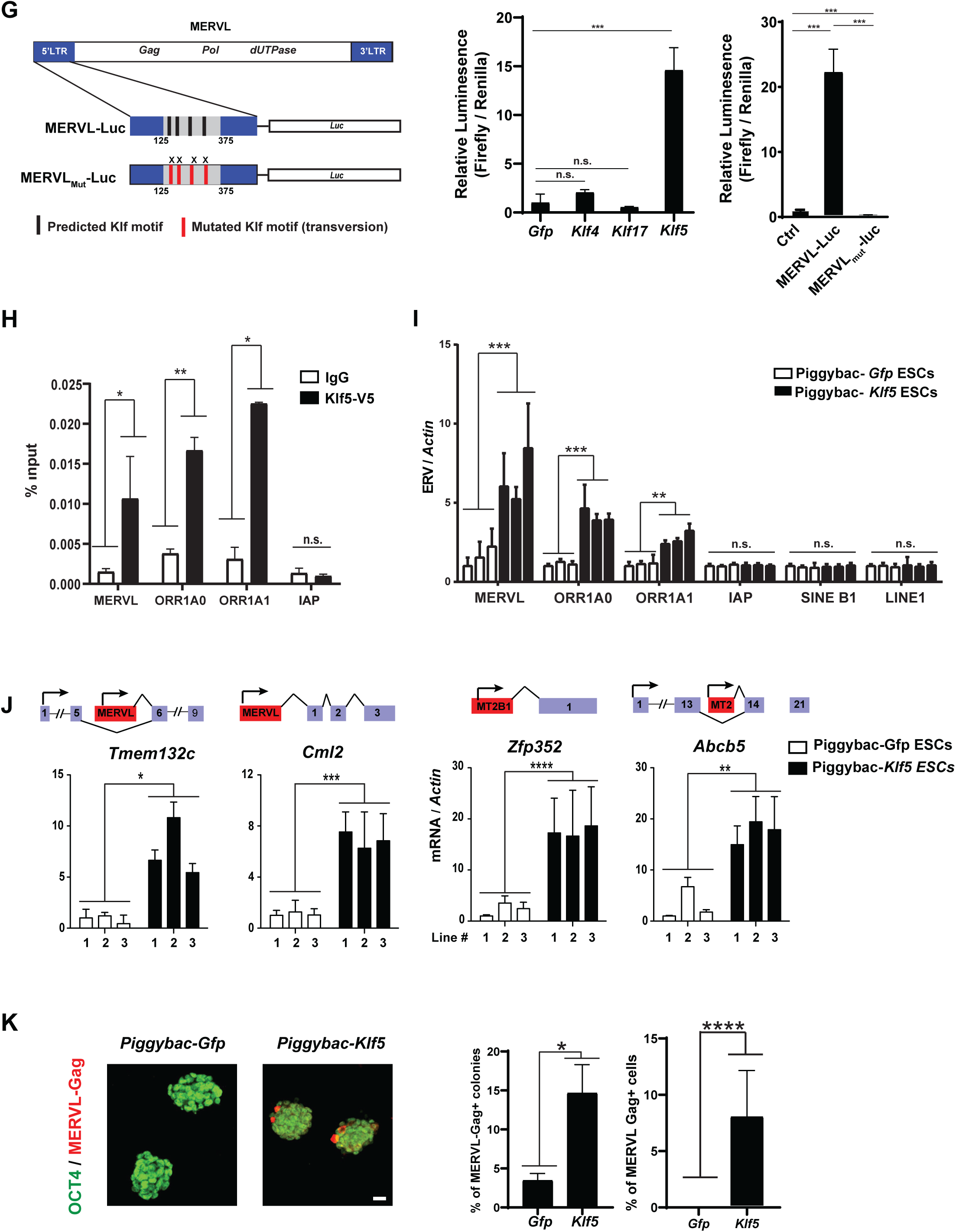
Klf5 directly regulates three major 2C-specific ERV families in preimplantation embryos. **A**. A flowchart illustrating the strategy to identify key transcription factors that induce 2C-specific ERVs. Retrotransposon profiling using reanalyzed published (GSE45719) RNA-seq data of mouse preimplantation embryos identifies 2C-specific ERVs(Deng et al., 2014). LTR sequences of 2C-specific ERVs were subjected to HOMER motif enrichment analyses to predict transcription factors that regulate these ERVs. **B, C.** MERVL, ORR1A0 and ORR1A1 are three major families of 2C-specific ERVs in preimplantation embryos. **B**. (Top) A heatmap illustrates the dynamics and stage-specific expression of a cohort of retrotransposons that peak in 2C stage in mouse preimplantation development. Red, annotated elements of major 2C-specific ERV families; Pro, pronucleus; 2C, two-cell stage; 4C, four-cell stage; 8C, eight-cell stage; 16C, sixteen-cell stage; BL, blastocyst. (Bottom) Gene structures were shown as diagrams for MERVL, ORR1A0 and ORR1A1. **C**. Single embryo real time PCR analyses using pronuclear (n= 4), 2C (n= 9), 8C (n= 3), and blastocyst embryos (n= 5) experimentally validated the 2C-specfic induction of MERVL (Pro vs 2C, \**P* = 0.0116, t = 3.024, df = 11; Pro vs 8C \*\*\*\**P* < 0.0001, t = 13.57, df = 5), ORR1A0(Pro vs 2C \**P* = 0.0295, t = 2.843, df = 6; Pro vs 8C \*\*\**P* = 0.0002, t = 9.278, df = 5) and ORR1A1(Pro vs 2C \*\**P* = 0.0063, t = 3.447, df = 10; Pro vs 8C \**P* = 0.0103, t = 4.567, df = 4). Error bars = s.d. **D**. MERVL, ORR1A0 and ORR1A1 are coordinately induced in multiple lines of 2C-like ESCs. (Left) Upon bioinformatic analyses of published RNA-seq data(Ishiuchi et al., 2015b), *Caf-1* deficient ESCs, including *P60* knockdown and *P150* knockdown ESCs, exhibited a coordinated induction of MERVL, ORR1A0 and ORR1A1 (MERVL, control vs *P60* knockdown, adjusted *P* = 1.99e-224, control vs *P150* knockdown, adjusted *P* =2.14e-212; ORR1A0, control vs *P60* knockdown adjusted *P* = 9.79e-64, control vs *P150* knockdown, adjusted *P* = 2.67e-86; ORR1A1, control vs *P60* knockdown, adjusted *P* = 1.92e-81, control vs *P150* knockdown, adjusted *P* = 2.68e-86). Error bars = s.d. P value was computed with the DESeq2 package in R. (Right) Real time PCR analyses confirmed a coordinated induction of ORR1A0, ORR1A1 and MERVL in *Lsd1* ^GT/GT^ ESCs. **E.** HOMER motif analyses predicted the enrichment of Klf binding motifs in the LTRs of 2C-specific ERVs. Binding motifs of Klf-family transcription factors are strongly enriched in LTRs of MERVL, ORR1A0 and ORR1A1, compared to scrambled control sequences. Dux and Gata2 are also labeled, highlighting the specific enrichment of their binding motifs in MERVL LTRs. **F**. *Klf5*, *Klf4* and *Klf17* are the major Klf transcription factors expressed in mouse early preimplantation embryos. Pseudotime expression plot showing the dynamic expression profiles of *Klf17, Klf4* and *Klf5* across preimplantation development. Klf5 exhibited an early induction that peaked at 4C stage, and its expression persisted throughout preimplantation development. **G**. Overexpression of *Klf5*, but not *Klf4* or *Klf17*, specifically induces the MERVL-luc reporter. (Left) A diagram showing the structure of the MERVL-luc and MERVL_Mut_-Luc reporters, which contained four predicted Klf binding motifs and four mutated motifs, respectively. The four predicted Klf binding motifs reside in the minimal MERVL LTR fragment (125bp – 375bp) required to recapitulate MERVL expression in ESCs(Choi et al., 2017; Macfarlan et al., 2011). (Right) overexpression of Klf5, but not Klf4 or Klf17, in HEK cells, along with a MERVL-Luc reporter, induced an increase in luciferase activity. Klf5, n=3, *P* = 0.0007, df = 4, t = 9.456; Klf4, n=2, n.s.; Klf17, n=2, n.s. Mutations of all four predicted Klf binding motifs in the MERVL_Mut_-Luc reporter abolished Klf5-dependent regulation. n = 3; error bars = s.d.; miniP-luc vs MERVL-luc, *P* = 0.0005, df = 4, t = 10.39; MERVL-luc vs MERVL_Mut_-Luc, *P* = 0.0004, df = 4, t = 0.73. **H-J.** *Klf5* overexpression in ESCs specifically induces MERVL, ORR1A0 and ORR1A1, and promotes a robust activation of the MERVL-associated transcriptome. **H.** Klf5 specifically occupies to the LTRs of major 2C ERVs. Chromatin immunoprecipitation (ChIP) of Klf5 in ESCs revealed specific Klf5 association with the LTRs of MERVL, ORR1A0 and ORR1A1. Two independent, passage matched ESC lines were tested. Error bars, s.d.; MERVL, \**P* = 0.0428, df = 2, t = 4.679, ORR1A0, \*\**P* = 0.0069, df = 2, t = 12.00; ORR1A1, \**P* = 0.0264, df = 2, t = 6.030. **I.** Three independent ESC lines were measured using real time PCR analyses. Error bars, s.d.; MERVL: \*\**P* = 0.0085, df = 4, t = 4.829; ORR1A0, \**P* = 0.0003, df = 4, t = 11.83; ORR1A1, \**P* = 0.0033, df = 4, t = 6.258 **J.** Specific MERVL elements can serve as alternative promoters to generate preimplantation-specific gene isoforms. MERVL-dependent gene isoforms were strongly induced in ESCs upon *Klf5* overexpression. *Zfp352, ****P* < 0.0001, df = 4, t = 16.12; *Abcb5*, \*\**P* = 0.0030, df = 4, t = 6.408; *Tmem132c*, \**P* = 0.0147, df = 4, t = 4.110; *Cml2,* \*\**P* = 0.0001, df = 4, t = 15.18. **K**. A subset of *Klf5-overexpressing* ESCs exhibited a strong expression of MERVL-Gag in immunofluorescence staining. The expression of MERVL-Gag in *Klf5-overexpressing* ESCs was mutually exclusive with that of Oct4 (Left). The percentage of MERVL-Gag positive ESC colonies (ESC colonies with 1 or more MERVL-Gag positive cells) and the percentage of MERVL-Gag positive cells in the total population were each quantified using fluorescent staining (Right). Scale bar, 20μm. error bars = s.d.. % of MERVL Gag positive colonies: \**P* = 0.0405, df = 4, t = 2.987. % of MERVL Gag positive cells: \*\*\*\**P* < 0.0001, df = 32, t = 6.712. * *P* < 0.05, ** *P* < 0.01, *** *P* < 0.001, **** *P* < 0.0001, *n.s*., not significant. All *P* values were computed using unpaired, two-tailed student’s t-test unless otherwise indicated.

The transcriptional regulatory sequences of ERVs are contained within their 5’ LTRs, often ∼300 - 500 bp in length (McCarthy and McDonald, 2004). Alignment of the consensus LTRs of the three 2C-specific ERVs revealed homology between ORR1A0 and ORR1A1 (96% identity), which shared 42% and 41% sequence identity, respectively, to MERVL LTR (Sup Fig. S1B). This finding suggests both coordinated and distinct transcriptional regulation of these 2C-specific ERVs.

### Identification of *Klf5* as a driver of 2C/4C-ERVs

To identify putative upstream regulators of this 2C cohort, we employed transcription factor motif enrichment analysis using all annotated LTR elements for each ERV family in the DFAM repository (Hubley et al., 2016), (Benner et al., 2017). A number of experimentally validated MERVL regulators emerged from these analyses, such as Dux, Gata2, Retinoic acid receptor alpha (Rara) Zfp281 and Tbx-family factors, indicating the power of this approach (Choi et al., 2017; Dai et al., 2017; Dan et al., 2013; Hendrickson et al., 2017; Tagliaferri et al., 2020). Interestingly, motifs for Klf transcription factors were the most enriched in the LTRs of MERVL, ORR1A0, and ORR1A1 (Fig. 1E, Sup Table S1), each of which harbors four predicted Klf binding motifs. In contrast, Dux and Gata2 exhibit a binding motif enrichment only within the LTRs of MERVL LTRs, but not ORR1A0/ORR1A1 (Fig. 1E, Sup Table S1). Consistently, in published ESC ChIP-seq data (Hendrickson et al., 2017), no direct Dux binding to ORR1A0 or ORR1A1 elements was detected (Sup Fig. S1C).

Krüppel-like factors (Klfs) are evolutionarily conserved zinc-finger transcription factors that have a pivotal role in embryonic development and pluripotent stem cells (Presnell et al., 2015). The mouse genome encodes 17 annotated Klf transcription factors, all of which possess a highly conserved C-terminal DNA binding domain that recognizes guanine-cytosine rich regions and CACCGT box motifs (Presnell et al., 2015). *Klf5*, *Klf4* and *Klf17* are the most highly expressed Klf transcription factors in mouse preimplantation embryos (Fig 1F, Sup Fig. S1D). While several Klf factors, including Klf4, Klf5 and Klf2, have been shown to function redundantly to sustain pluripotency in ESCs(Yamane et al., 2018), Klf5 is the only factor whose deficiency in mouse embryos impairs preimplantation cell fate decisions (Azami et al., 2017; Ema et al., 2008; Lin et al., 2010; Presnell et al., 2015). In comparison, *Klf4* or *Klf2* individual knockout yield no obvious preimplantation defects (Katz et al., 2002)(Wani et al., 1998).

To determine the Klf(s) capable of directly regulating 2C-specific ERVs, and possibly, the bi-potent cell fate, we constructed luciferase reporters driven by the MERVL LTR (MERVL-Luc), which faithfully recapitulates MERVL expression (Choi et al., 2017; Macfarlan et al., 2011). While Klf5 was able to activate MERVL-luc in HEK cells, neither Klf4 nor Klf17 had a similar effect (Fig. 1G). The MERVL LTR contains 4 predicted Klf5 binding motifs, and transversion mutations of these sites abolished Klf5-dependent luciferase induction (Fig. 1G). To ascertain whether Klf5 directly mediated MERVL-luc induction, we performed chromatin immunoprecipitation (ChIP) using *Klf5-overexpressing* ESCs and confirmed specific Klf5 occupancy on the LTRs of MERVL, ORR1A0, and ORR1A1 (Fig. 1H). Consistently, *Klf5* overexpression specifically upregulated MERVL, ORR1A0, and ORR1A1, along with 2C-specific, MERVL driven 2C gene isoforms (Fig. 1I, 1J). No induction of other retrotransposon families, such as IAP, LINE1, and SINE B1, was observed (Fig. 1I). *Klf5* overexpression in ESCs also activated a tdtomato reporter driven by the MERVL LTR (Sup Fig. S1F). MERVL induction by Klf5 in ESCs was further confirmed by immunostaining for MERVL-Gag protein, with ∼15% ESC colonies containing MERVL Gag-positive cells and 5-8% MERVL Gag-positive cells overall (Fig. 1K). In comparison, control ESCs contained <0.1% MERVL+ cells (Fig 1K), indicating that Klf5 overexpression shifted the equilibrium in ESC culture in favor of metastable MERVL^+^ cells. Similar to other reported MERVL^+^ ESCs(Choi et al., 2017; Hendrickson et al., 2017; Macfarlan et al., 2012), overexpression of *Klf5* in ESCs activated MERVL at the expense of Oct4 protein expression, as MERVL-Gag and Oct expression was mutually exclusive (Fig. 1K). Importantly, *Klf5* overexpression levels in ESCs were comparable to the level of *Klf5* in blastocysts, within a physiologically relevant range (Sup Fig. S1G), and RNA-seq in *Klf5* overexpressing ESCs further confirmed the ability of Klf5 to upregulate 2C-specific genes (Sup Fig. S1H). Altogether, our data establish Klf5 as a direct regulator of all three 2C specific ERVs, including MERVL, ORR1A0 and ORR1A1.

### *Klf5* confers bi-potential cell fate *in vitro* and *in vivo*

Since Klf5 induces MERVL, ORR1A0, and ORR1A1 ERVs, which collectively marked bi-potent blastomeres and ESC populations (Fig. 1B-1D), we hypothesized that *Klf5* could functionally confer bi-potential cell fate in ESCs. In line with this hypothesis, teratomas generated from *Klf5* overexpressing ESCs contained cells expressing markers of TE (*Cdx2* and *Elf5*), primitive endoderm (PrE) (*Gata4*, *Gata6* and *Sox17*), and all three embryonic germ layers (*Pax6*, *Brachyury* and *Foxa2*) (Fig. 2A, 2B, Sup Fig. S2A, S2B). In comparison, control teratomas only induced molecular markers of embryonic lineages (Fig. 2B). In particular, we identified teratoma cells reminiscent of placental trophoblast giant cells, with strong PL-1 (placental lactogen 1) expression, large cell volume, enlarged nuclei, and proximity to internal hemorrhages (Fig 2A). Similarly, embryoid bodies (EBs) generated from *Klf5-overexpressing* ESCs, but not control ESCs, showed induction of extra-embryonic markers of the TE, PrE and placental trophoblast lineages (Fig 2C, Sup Fig. S2C)

**Figure 2.**
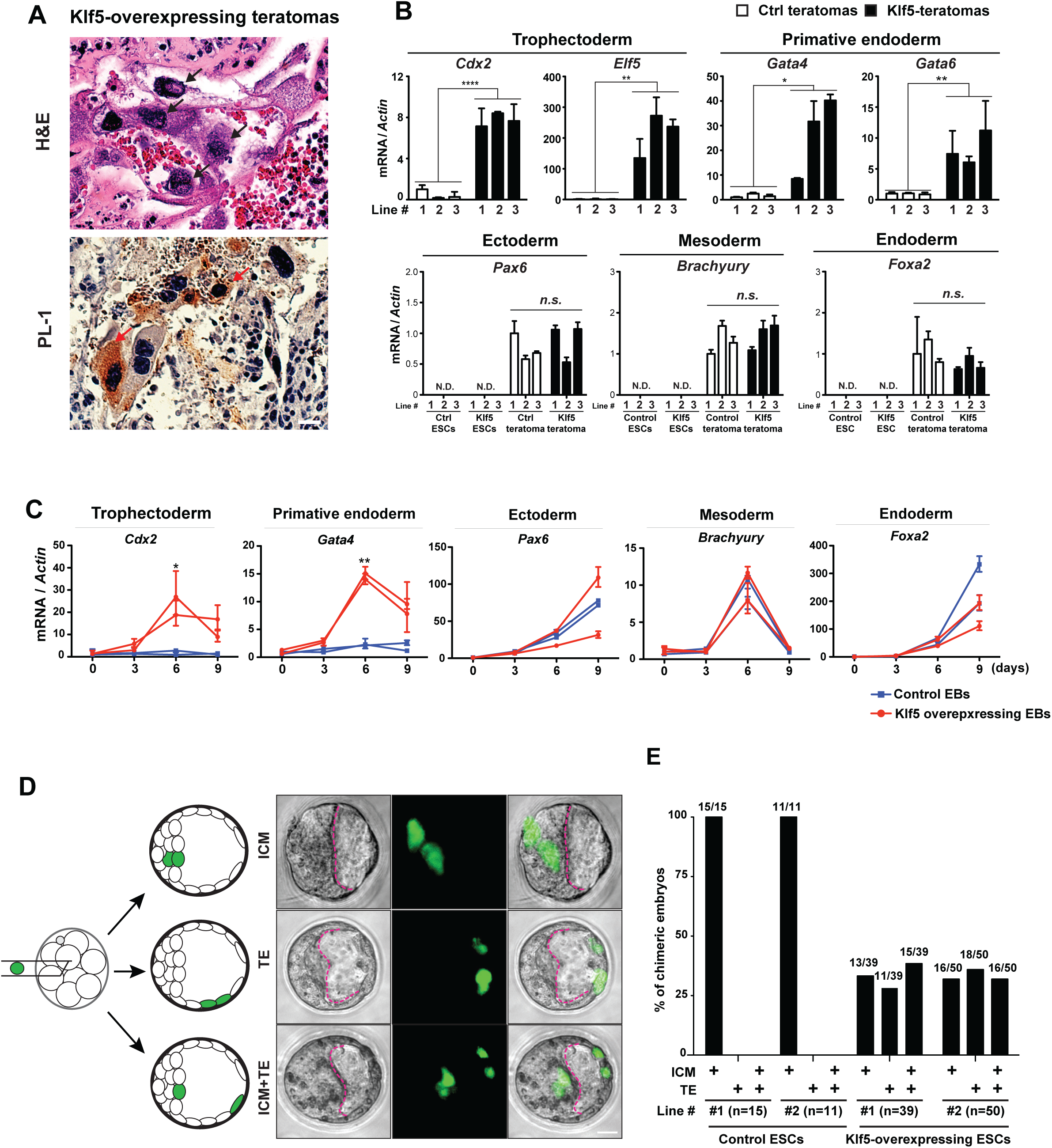
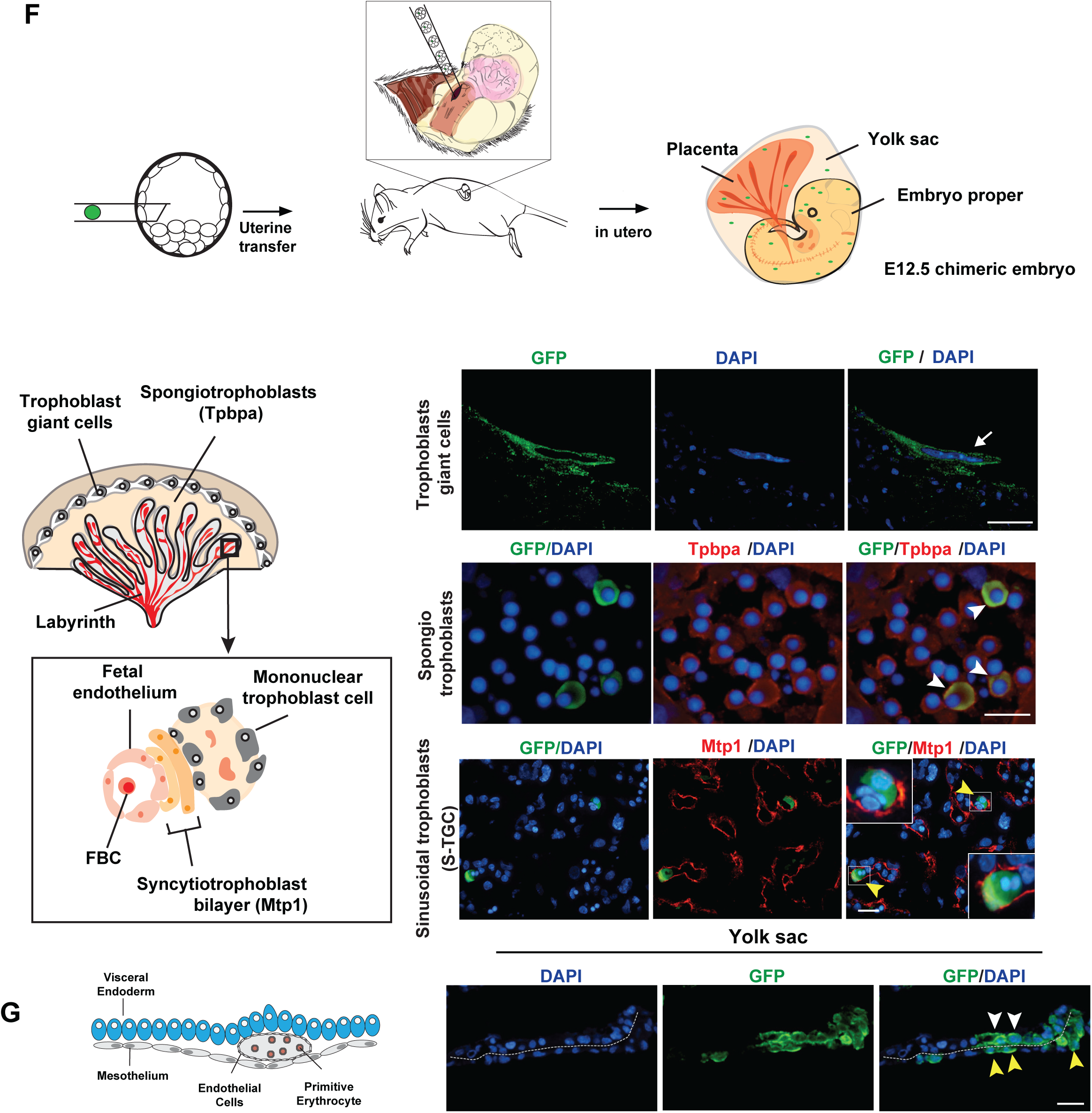
Klf5 overexpression confers a bi-potential cell fate in ESCs both *in vitro* and *in vivo*. **A.** Teratomas derived from *Klf5-overexpressing* ESCs contain embryonic and extra-embryonic cell lineages. Teratomas generated from *Klf5-overexpressing* ESCs contain cells with characteristic placental trophoblast giant cell like morphology (black arrows, top) and strong placental lactogen 1 (PL-1) expression (red arrows, bottom). Scale bars, 50 μm. **B.** The TE markers (*Cdx2* and *Elf5*) and the primitive endoderm (PE) markers (*Gata4* and *Gata6*) were highly induced in *Klf5-overexpressing* teratomas in real-time PCR analyses, but not in control teratomas. In contrast, the expression of *Pax6* (an ectoderm marker), *Brachyury* (a mesoderm marker) and *Foxa2* (an endoderm marker) are similarly induced in control and *Klf5-overexpressing* teratomas. Teratomas were generated from three independent pairs of passage-controlled control and *Klf5-overexpressing* ESC lines. Error bars: *s.d*.; *Cdx2*, **** *P* < 0.0001, df = 4, t = 16.05; *Elf5*, ** *P* = 0.0066, df = 4, t = 5.182; *Gata4*, \**P* = 0.0427, df = 4, t = 2.934; *Gata6*, ** *P* = 0.0092, df = 4, t = 4.713. **C**. *Klf5-overexpressing* EBs, but not control EBs, showed a significant induction in TE markers *Cdx2* and PE markers *Gata4*. In contrast, they similarly induced markers of all three germ layers, *Pax6*, *Brachyury* and *Foxa2.* EBs were generated from two independent pairs of passage-controlled control and *Klf5-overexpressing* ESC lines. Error bars, *s.d.*; *Cdx2* (day 6), * *P* = 0.0372, df = 2, t = 5.036; *Gata4* (day 6), ** *P* = 0.0019, df = 2, t = 23.20. **D, E.** Single *Klf5-overexpressing* ESCs confer bi-potential cell fate in chimeric blastocyst embryos. Representative images (**D**) and quantitation (**E**) of chimeric blastocyst embryos with ESC contribution to the ICM, the TE, or both. **D.** A diagram illustrates the experimental strategy to generate chimeric blastocysts embryos by microinjecting a single GFP-labeled, *Klf5-overexpressing* ESC into each C57BL/6N recipient morula. ESC contribution to the ICM and/or the TE was determined by the localization of GFP-positive ESC progenies in fluorescence imaging. Scale bar, 20 μm. **D, E.** Chimeric blastocysts were generated from two independent pairs of passage-controlled control and *Klf5-overexpressing* ESCs lines. **F, G**. Single *Klf5-overexpressing* ESCs confer bi-potential cell fate in chimeric E12.5 embryos, generating terminally differentiated extra-embryonic placental and yolk sac lineages. **F**. (Top) A diagram illustrating the experimental strategy to generate chimeric E12.5 embryos by microinjecting a single GFP-labeled, *Klf5-overexpressing* ESC into each C57BL/6N recipient blastocysts, followed by embryo transfer into pseudo-pregnant mother. (Bottom left) A diagram illustrating several terminally differentiated cell lineages of placenta. (Bottom right) In E12.5 chimeric embryos, progenies from a single GFP-labeled, *Klf5* overexpressed ESC generated trophoblast giant cells with a characteristic cellular morphology (white arrows), spongiotrophoblasts with Tpbpa expression (white arrowheads), and syncytio trophoblasts with Mtp1 expression (yellow arrowheads). Scale bars, 100μm. **G**. (Left) A diagram illustrating several terminally differentiated cell lineages of yolk sac. (Right) A single GFP-labeled, *Klf5-overexpressing* ESC yield terminally differentiated visceral endoderm cells (white arrowheads) and embryonic mesothelium (yellow arrowheads) in yolk sac of E12.5 chimeric embryos. Extra-embryonic visceral endoderm cells were identified based on the bilaminar structure of the yolk sac and their characteristic columnar epithelial morphology. Scale bar, 20 μm; * *P* < 0.05, ** *P* < 0.01, *** *P* < 0.001, *n.s*., not significant. All *P* values were computed using unpaired, two-tailed student’s t-test.

In teratomas and EBs, multiple MERVL+ cells collectively contribute to both embryonic and extraembryonic cell types, making it unclear if MERVL+ cells have *bona fide* bi-potential cell fate. The bi-potential cell fate of early blastomeres is strictly defined by the capability of a single cell to contribute to both embryonic and extraembryonic lineages (Wigger et al., 2017), (Wu et al., 2017). Hence, we microinjected single, GFP-labeled control or *Klf5-overexpressing* ESCs into each C57BL/6J recipient 8C embryos and analyzed the resulting chimeric blastocysts (Fig. 2D). While control ESCs invariably contributed to the ICM of chimeric blastocysts (Sup Fig. S2D), individual *Klf5-overexpressing* ESCs colonized the ICM, TE, or both in chimeric embryos (Fig. 2D, 2E). Particularly, nearly a third of chimeric blastocysts contained GFP positive progenies from a single *Klf5-overexpressing* ESC in both ICM and TE (Fig. 2D, 2E). Intriguingly, the GFP+ ICM cells derived from *Klf5*-expressing ESCs strongly express Nanog in chimeric blastocysts, similar to their neighboring ICM cells. Yet the GFP+ TE cells exhibit an intermediate phenotype: They localized to the TE compartment, took on a typical TE morphology, lost Nanog expression, but failed to robustly express Cdx2 as their neighboring TE cells (Sup Fig. S2E). This is likely due to the delayed kinetics of *Cdx2* activation upon Klf5-overexpressing ESCs withdrawal from its pluripotent cell fate to take a TE cell fate. These TE cells derived from *Klf5*-overexpressing ESCs, while acquired extra-embryonic potency, have a reduced efficiency for extra-embryonic differentiation compared to normal TE cells.

We next generated E12.5 chimeric embryos by injecting single *Klf5-overexpressing* ESCs into recipient blastocysts. These E12.5 chimeric embryos contained ESC-derived cell lineages in both the embryo proper and extra-embryonic placental and yolk sac lineages (Fig. 2F-2H, Sup Fig. S2F, Sup Table S2). *Klf5-overexpressing* ESCs gave rise to terminally differentiated trophoblast giant cells, spongiotrophoblasts of the placenta, as well as the yolk sac visceral endoderm (Fig. 2F, 2G). In addition, GFP positive cells with large cytoplasmic to nuclear ratio were found proximal to Mtp1 positive syncytiotrophoblast II (SynII) cells, which morphologically resemble sinusoidal trophoblast giant cells (s-TGCs) (Fig. 2F). Our findings suggest that, in each E12.5 chimeric embryo, a single injected *Klf5-overexpressing* ESC underwent substantial proliferation prior to terminal lineage commitment during normal development. E12.5 chimeric embryos generated by injecting 10-15 *Klf5-overexpressing* ESCs yielded similar results, generating terminally differentiated embryonic lineages, as well as extra-embryonic placental and yolk sac lineages (Sup Fig. S2G-S2H). In all our Klf5 overexpression studies, *Klf5* overexpressing ESCs have a level of *Klf5* expression comparable to that of blastocysts (Sup Fig. S1G), suggesting that the *Klf5* overexpression phenotype we observed in ESCs is caused by a physiologically relevant level of *Klf5* expression. The bi-potential phenotype of *Klf5* over-expressing ESCs documented here is among the best documented, *in vivo* bi-potential phenotype generated by a single cell, suggesting *Klf5* as an important bi-potential regulator.

### *Klf5* regulates both ICM and TE specification genes

To understand the molecular basis of Klf5 induced bi-potential cell fate in ESCs, we performed ChIP-seq using *Klf5-overexpressing* ESCs, and applied Ingenuity Pathway Analysis (IPA) on Klf5-bound genes (Fig. 3A). Interestingly, Klf5 occupancies were significantly enriched for genes regulating ESC pluripotency, Hippo signaling and blastocyst development (Fig. 3A, Sup Table S3), many of which promoted cell fate specification of the ICM or TE and displayed dynamic expression in cleavage stage embryos (Fig. 3B). Specifically, ICM-specific transcription factors (*Nanog* and *Klf4*) and TE-specific transcription factors (*Tead4* and *Cdx2*) all exhibited Klf5 binding in their putative promoter / enhancer regions (Fig. 3C); the corresponding Klf5 peaks invariably harbored multiple predicted Klf binding motifs (Sup Fig S3A). Additionally, Klf5 peaks were also observed in 2C-ERV LTRs that drove 2C-specific gene isoforms (Sup Fig. S3B).

**Figure 3.**
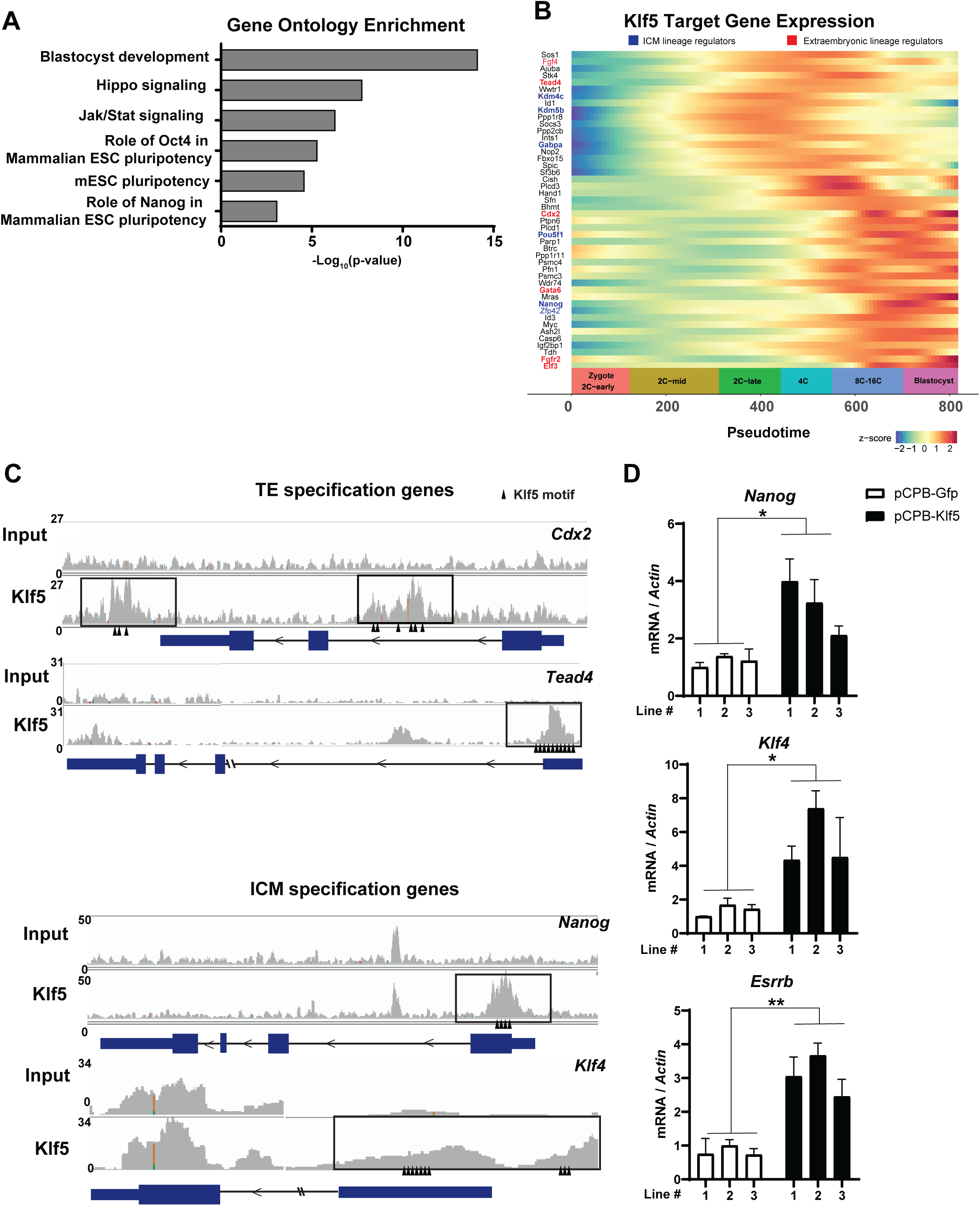
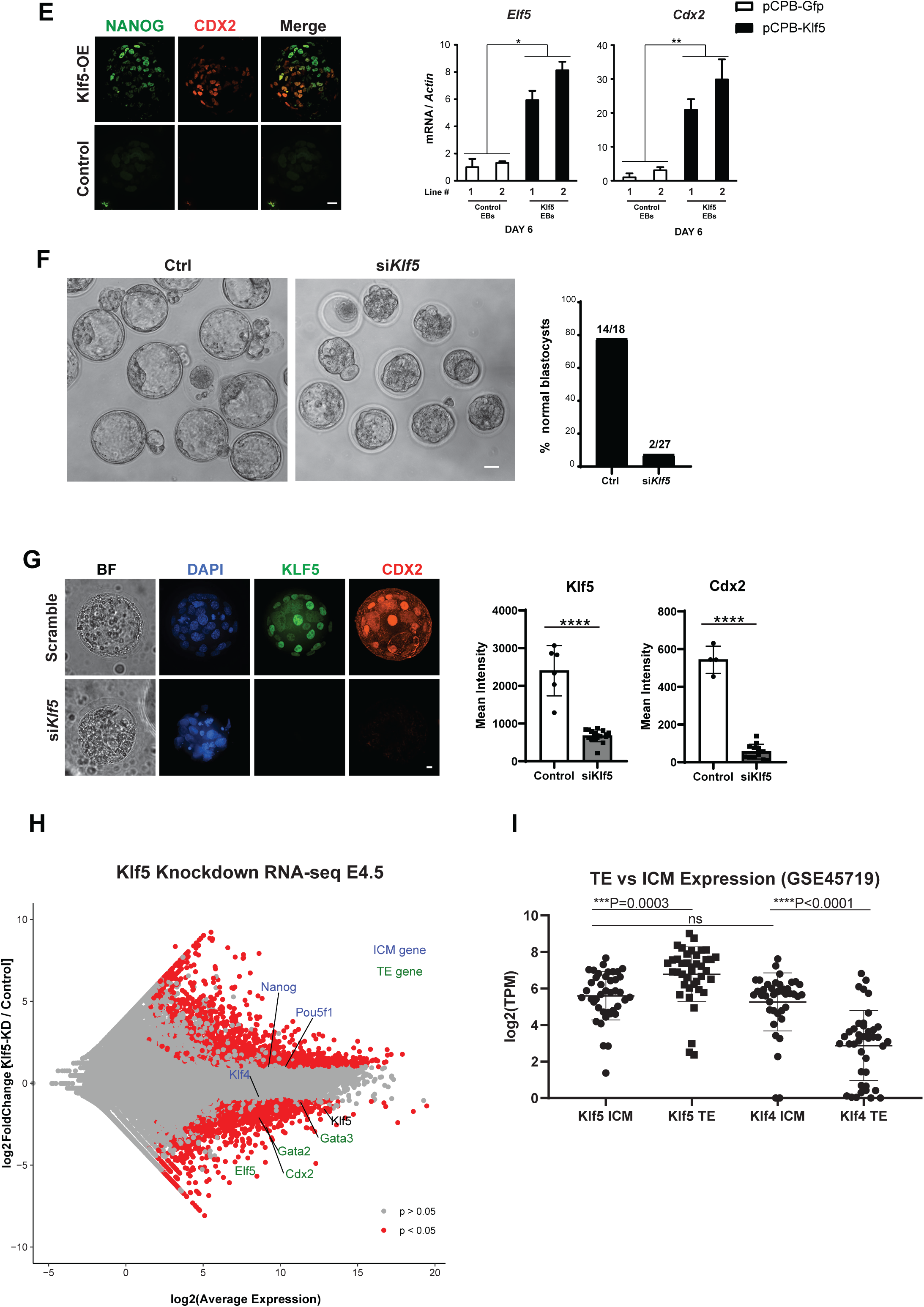
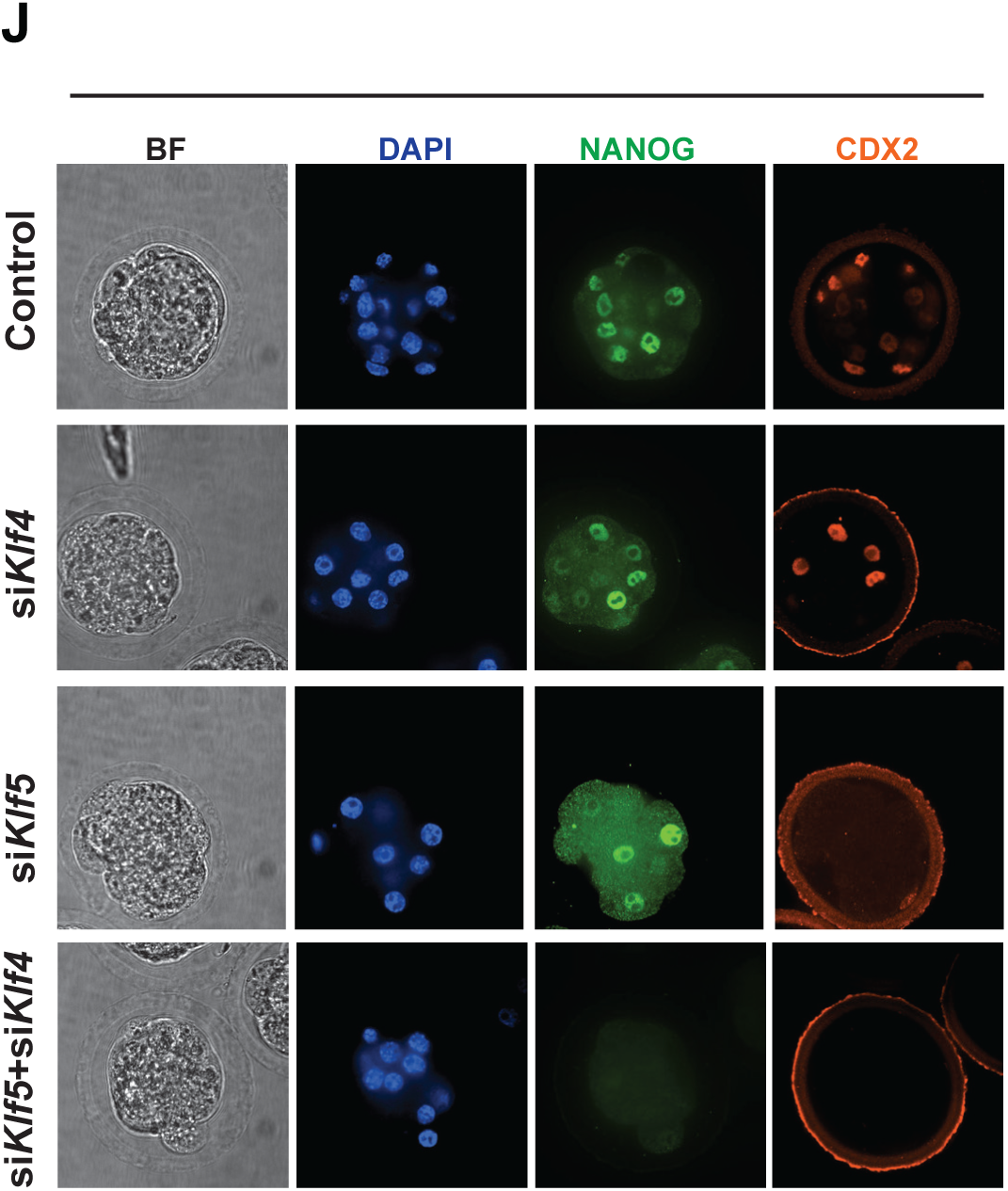
Klf5 promotes dual induction of cell fate specification genes for both embryonic and extraembryonic commitment. **A.** Ingenuity pathway analysis reveals the enriched functional terms among Klf5 ChIP-seq targets. **B.** A heatmap illustrates the expression profiles of the 70 most dynamically expressed *Klf5* targets in preimplantation development, which include multiple ICM (blue) and TE (red) specification genes. **C.** Read density of Klf5 ChIP-seq reads illustrates the enrichment of Klf5 occupancy on the cis-regulatory elements of lineage specification genes, including TE specification genes (*Cdx2* and *Tead4*), as well as ICM specification genes (*Nanog* and *Klf4)*. **D.** *Klf5* overexpression in ESCs elevates the expression of pluripotency-specific transcription factors, including *Nanog*, *Klf4* and *Errsb*. Nanog: \**P=* 0.0264, df = 4, t = 3.436, Klf4: \**P=* 0.0162, df = 4, t = 3.992, Essrb: \*\**P=* 0.0035, df = 4, t = 6.153. N = 3 passage-controlled ESC lines. Error bars = s.d. **E.** EBs derived from *Klf5-overexpressing* ESCs contain Cdx2-positive cells as well as Nanog-positive cells, whereas control EBs failed to activate Cdx2 and completely lost Nanog expression. Real time PCR analyses confirmed the induction of TE markers (Cdx2 and Elf5) in *Klf5-overexpressing* EBs. Scale bars, 100 μm (left). Error bars, s.d..; *Cdx2*, \*\**P* = 0.0372, df = 2, t = 5.036; *Elf5*, \**P* = 0.0339, df = 2, t = 5.289. **F, G.** *Klf5* knockdown stalls preimplantation development before the blastocyst stage and abolishes Cdx2 expression in TE. **F.** Representative bright field images (left) and quantitation (right) are shown for E4.0 *Klf5* knockdown (siKlf5) or control (scramble siRNA) blastocyst embryos. **G**. Representative Cdx2 immunofluorescence images (left) and relative fluorescence quantitation (right) are shown for control and *Klf5* knockdown blastocyst embryos. 3 independent *Klf5* knockdown experiments were performed using a total of 18 control embryos and 27 *Klf5* knockdown embryos. Scale bar = 20uM; Klf5, \*\*\*\**P*= < 0.0001, df = 22, t = 10.62; Cdx2, \*\*\*\**P* = < 0.0001, df = 14, t = 17.46. **H**. MA-plot of RNA-seq data from control versus *Klf5* knockdown E4.5 embryos confirming that *Klf5* depletion leads to defects in TE specification as shown by reduced expression of TE-marker genes: *Cdx2, Elf5, Gata2* and *Gata3* whereas expression of ICM genes *Klf4, Nanog* and *Pou5f1* is not significantly impacted. **I.** Expression analysis of *Klf4* and *Klf5* in TE and ICM cells demonstrating that *Klf5* mRNA is enriched in the TE whereas *Klf4* is expressed at comparable levels to *Klf5* in the ICM and is expressed at a significantly lower level in the TE. **J.** Immunofluorescent panel from control, *Klf4*, *Klf5* and *Klf4 + Klf5* knockdown embryos at E3.25 stained for NANOG and CDX2 protein demonstrating that Klf4 and Klf5 cooperate during ICM specification. Scale bar = 20uM.

While Klf5 has the ability to induce both ICM and TE genes based on our ChIP-seq data, Klf5 mediates gene transcription in a cell type- and context-specific manner. In pluripotent ESCs, *Klf5* overexpression induced multiple pluripotency transcription factors, including *Nanog*, *Klf4* and *Esrrb* (Fig. 3D), but had little effect on TE genes, possibly due to strong epigenetic silencing of TE genes in standard ESC cultures (Niwa et al., 2005). In differentiating EBs derived from *Klf5-overexpressing* ESCs, we observed distinct Nanog^+^ or Cdx2^+^ cell populations, which were absent in control EBs (Fig. 3E). The strong induction of extra-embryonic TE markers, *Cdx2* and *Elf5,* was confirmed by real time PCR (Fig. 3E). Hence, *Klf5-overexpressing* ESCs yielded progenies expressing either ICM or TE genes within an EB (Fig. 3E), consistent with the bi-potential developmental plasticity of Klf5-expressing ESCs.

Prior to cell fate specification during the morula to blastocyst transition, modest co-expression of ICM and TE-specific transcription factors in early blastomeres is essential for establishing a bi-potential cell fate (Hirate et al., 2013; Korotkevich et al., 2017; Strumpf, 2005), (Pfeffer, 2018). These early blastomeres provide a unique cellular context where ICM and TE genes can be co-expressed at a modest level. The ability of Klf5 for dual induction of ICM and TE specification genes likely plays a role in conferring a cell state that allows differentiation into either ICM or TE lineages.

*Klf5* expression initiates in bi-potent blastomeres of cleavage stage embryos (Fig. 1F). Subsequently, Klf5 exhibits both ICM and TE expression in blastocysts, albeit with a stronger TE enrichment (Sup Fig. S3C). To investigate the role of *Klf5* in cell fate potency during preimplantation development, we efficiently knocked down *Klf5* by 90% using RNA interference (RNAi) in zygotes (Sup Fig. S3D). Morphologically, *Klf5* knockdown embryos appeared normal until the morula stage; obvious defects arose with high penetrance during blastocoel formation. By E4.5, control embryos had typical blastocyst morphology, whereas *Klf5* knockdown embryos failed to robustly form a blastocoel cavity (Fig 3F). Immunofluorescence staining revealed a marked reduction of Cdx2 in E4.5 *Klf5* knockdown embryos (Fig 3G), consistent with a significant decrease in major TE specification genes (*Cdx2*, *Elf5* and *Tead4*) and placenta early development gene (*Esrrb*) as shown by real time PCR in control versus knockdown embryos (Sup Fig. S3D). In contrast, *Klf5* knock down alone failed to affect ICM specification genes or pluripotency genes *in vivo* (*Sox2*, *Oct4* and *Nanog*) (Sup Fig. S3D). RNA-seq in control and *Klf5* knockdown embryos further confirmed the different effects of *Klf5* on TE versus ICM *in vivo*. *Klf5* knockdown significantly decreased the level of TE specification genes in E4.5 blastocysts, while leaving the ICM specification genes largely unperturbed (Fig 3H).

To confirm these findings, we generated blastocyst embryos with a complete *Klf5* disruption, using CRISPR-mediated *Klf5* editing (Sup Fig. S3E). In *Klf5* knockout embryos, we detected a significant loss of Cdx2, but Oct4 expression was relatively intact (Sup Fig S3E). Hence, while *Klf5* has the ability to induce both ICM and TE specification genes, *Klf5* deficiency preferentially impairs TE cell fate, consistent with a strong Klf5 enrichment in the TE compartment (Sup Fig. S3C). Our findings seemingly differ from a previous study, which described impaired ICM and TE specification caused by *Klf5* deficiency(Lin et al., 2010). Nevertheless, a closer examination of their data indicates that the TE defects are the predominant preimplantation phenotype in *Klf5* knockout embryos, and the ICM defects are rather mild with incomplete penetrance (Azami et al., 2017; Lin et al., 2010). Hence, while Klf5 can induce both ICM and TE genes, it is essential for TE specification, but not for ICM specification.

Given the ability of *Klf5* to co-induce ICM and TE genes, the lack of an obvious ICM defect in Klf5 deficient embryos suggests that additional Klf transcription factor(s) could act redundantly to specify ICM. Interestingly, Klf5 is expressed in both ICM and TE, with a strong TE enrichment; *Klf4* is specifically enriched in the ICM, and its ICM expression level is comparable to that of *Klf5* (Fig. 3I). Consistent with this expression pattern, *Klf4* knockout in mice exhibits no obvious defects in preimplantation development(Katz et al., 2002), yet knocking down both *Klf5* and *Klf4* impairs both ICM and TE cell fate during the morula to blastocyst transition, as shown by defective Nanog and Cdx2 expression (Fig. 3J, Sup Fig. S3F). Interestingly, *Klf5* knockdown failed to alter Yap-1 staining in embryos, suggesting that the Hippo signaling is not regulated by *Klf5* during preimplantation development (Sup Fig. S3G). This is consistent with Klf5 enabling cell fate potency for both the ICM and the TE lineages, rather than specifying the TE lineages. Altogether, Klf transcription factors constitute a functional robust transcriptional network for bi-potential cell fate, with Klf5 being an essential regulator for enabling TE specification, and Klf4 and Klf5 being functionally redundant to enable ICM specification.

Interestingly, the ICM expression of Klf5 is observed in the epiblast as well as the PrE lineage in E4.0 blastocysts, as Klf5 and Sox17 are co-expressed in cells of the PrE compartment (Sup Fig. 3H). Real time PCR analyses of *Klf5* knockdown embryos also exhibited a marked downregulation of PrE specification genes *Gata6*, *Gata4* and *Sox17* (Sup Fig 3H). While this is consistent with the induction of PrE gene markers in *Klf5*-overexpressing EBs and teratomas (Fig. 2B), the decreased PrE gene expression in *Klf5* knockdown embryos can also be caused by their developmental arrest/delay. Taken together, our data are consistent with Klf5 acting upstream of the induction of PrE genes. Our findings contrast a previous study, which reported Klf5 as a suppressor for PrE specification (Azami et al., 2017). This study showed an increased percentage of PrE cells in *Klf5* knockout blastocyst embryos. Yet the significant decrease of the total cell number and the developmental arrest of *Klf5* knockout blastocysts have complicated the interpretation of this result, making it difficult to conclude Klf5 as a suppressor of the PrE cell fate(Azami et al., 2017). In addition, the similar expression of *Klf5* in PrE and epiblast cells is inconsistent with a suppressive role of *Klf5* in the PrE cell fate decision. It is clear that the strongest and the most direct phenotype of Klf5 deficient embryos is the impaired TE specification.

### *Klf5* is regulated by multiple 2C-transcription factors

Using all ESC ChIP-seq data in the Cistrome database (Cistrome DB) (Mei et al., 2017), we identified a number of transcription regulators with enriched occupancy proximal to *Klf5* (Fig. 4A, Sup Table S4). These putative Klf5 regulators were subjected to Ingenuity Pathway Analysis (IPA), and an enrichment emerged for genes regulating stem cell biology and preimplantation development (Sup Fig. S4A). In particular, multiple 2C-specific transcription factors, including Dux, Dppa2 and Tbx3, bound to putative Klf5 regulatory regions, either a region immediately upstream of the transcription start site (TSS), or a region within the intron 1 (Fig. 4A). In particular, Dux, the key regulator for 2C-like transcriptome (Eckersley-Maslin et al., 2019; Hendrickson et al., 2017; Hu et al., 2020; Iaco et al., 2017; Ishiuchi et al., 2015a), showed a strong ChIP-seq peak in *Klf5* intron 1 (Fig. 4A). Using published RNA-seq data on *Dux*-overexpressing ESCs (Hendrickson et al., 2017), we showed that *Dux* overexpression upregulates *Klf5* (Fig. 4B). In addition to *Dux*, *Dppa2* also induced *Klf5* expression and MERVL expression in ESCs (Fig. 4B) and this induction was independent of Dux. Finally, *Tbx3* overexpression in ESCs also induced *Klf5* (Fig. 4B). Hence, multiple 2C-specific transcription factors converge their direct regulation on the induction of *Klf5*.

**Fig. 4.**
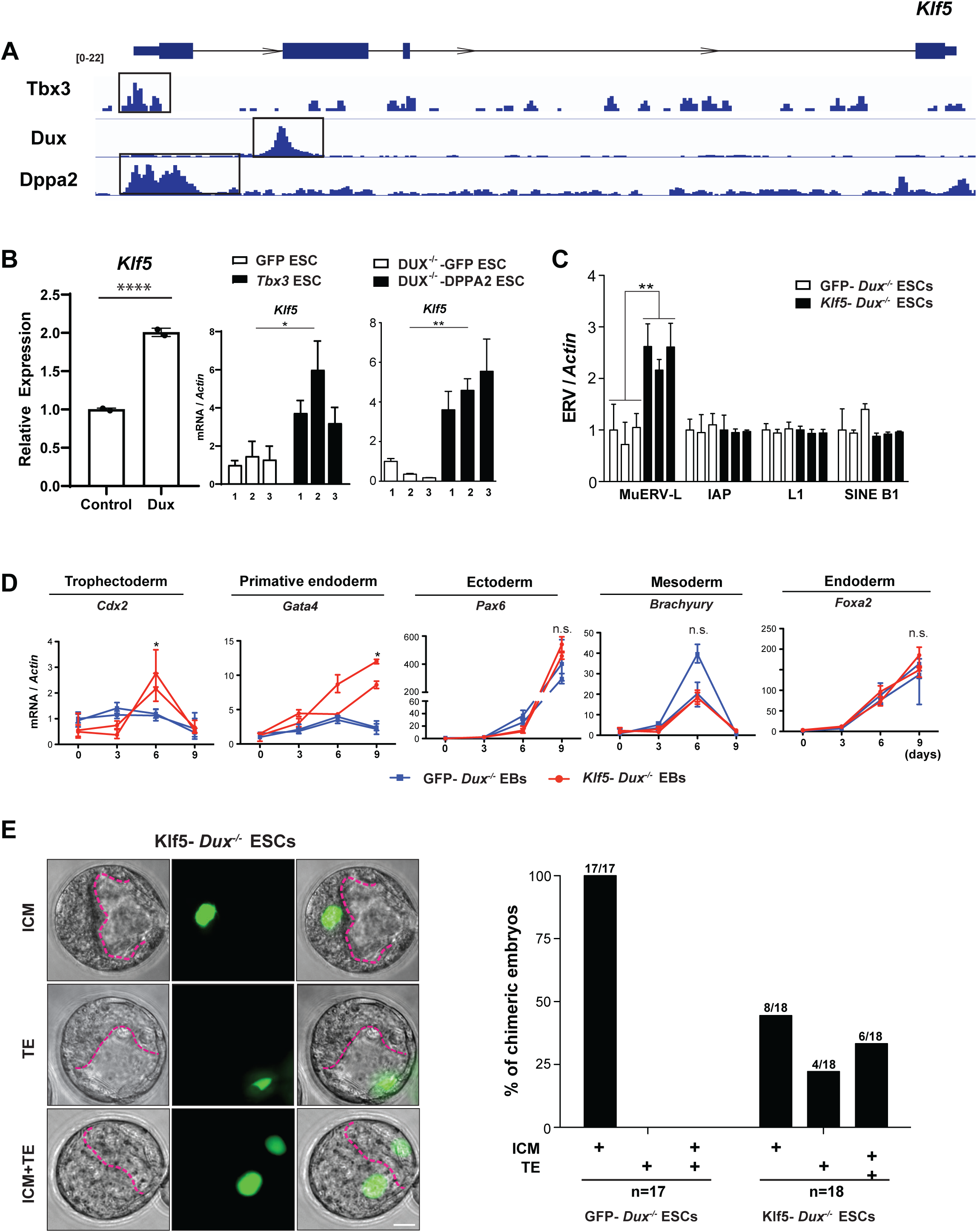
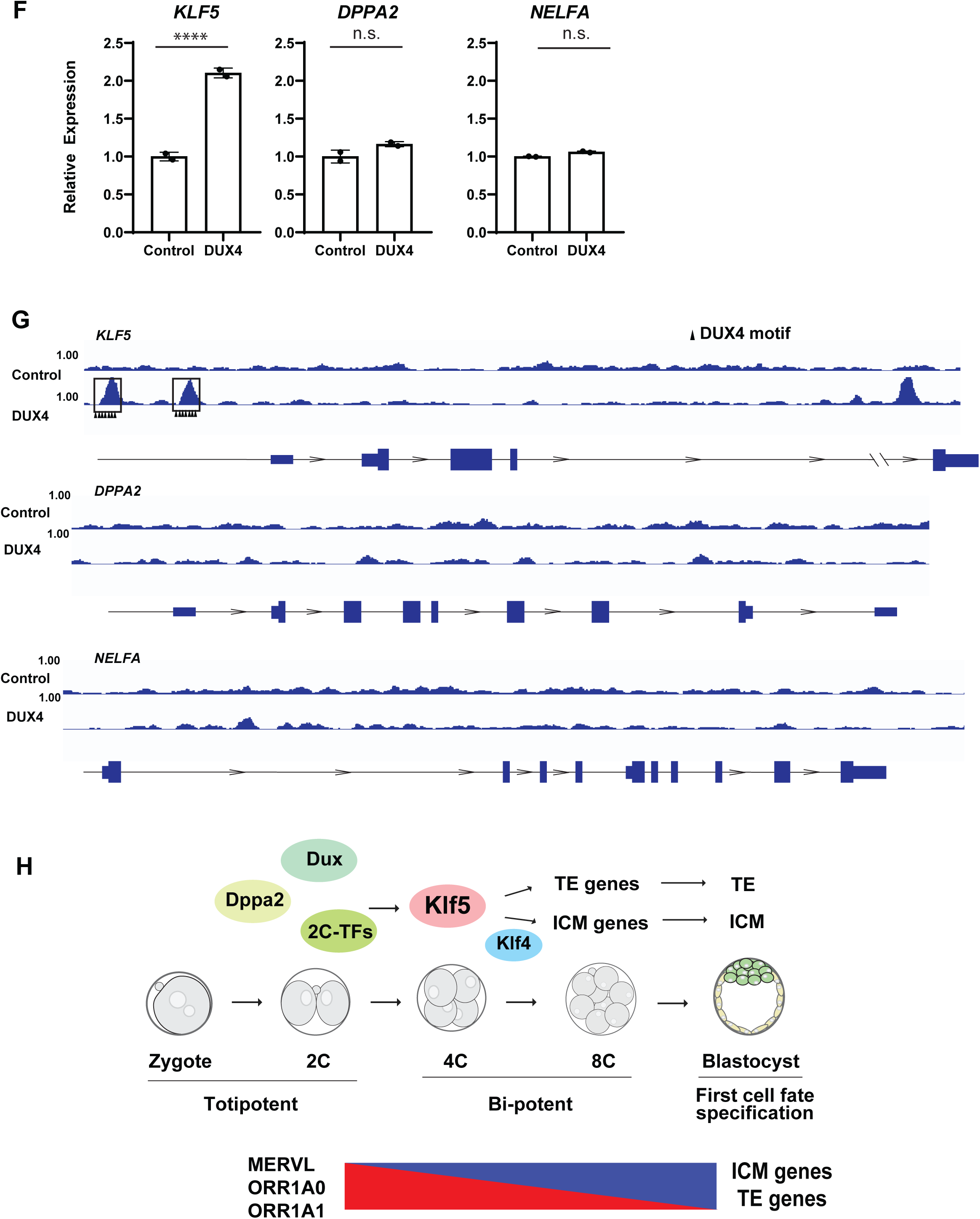
Klf5 induction is regulated by the 2C-specific transcription factor Dux. **A.** Multiple 2C-specific transcription factors show enrichment of occupancies proximal to *Klf5*. We mined all ESC ChIP-seq experiments in Cistrome DB and identified specific binding of Klf5 to a number of published 2C-specific transcription factors. The Y axis shows the coverage depth of ChIP-seq reads, ranging from 0 to 22 for each transcription factor. **B** *Dux, Tbx3,* and *Dppa2* overexpression in ESCs induces *Klf5*. Using RNA-seq data from *Dux*-overexpressing and control ESCs(Hendrickson et al., 2017), we demonstrated a robust induction of *Klf5* by *Dux*. Additionally, *Tbx3* and *Dppa2* overexpression also upregulates *Klf5*. Error bars = s.d.; *Klf5* (Dux overexpression), *P* < 0.0001; *Klf5(Tbx3* overexpression) *P =* 0.0244, df = 4, t = 3.522; *Klf5(Dppa2* overexpression) \*\**P* = 0.0027, df = 4, t = 6.622. *P* values for Dux overexpression were calculated with the DEseq2 package in R, otherwise *P* values were calculated on the basis of an unpaired Student’s t-test. **C**. *Klf5* acts downstream of *Dux* to induce MERVL. Real time PCR analyses detected specific MERVL induction following Klf5 overexpression in a *Dux* knockout ESCs. MERVL, \*\**P* = 0.0011, df = 4, t = 8.407. **D.** EBs derived from *Klf5-overexpressing Dux^-/-^* ESCs, but not control ESCs, showed significant induction of extra-embryonic TE marker *Cdx2* and primitive endoderm marker *Gata4*. Markers for endoderm, mesoderm and ectoderm (*Foxa2, Brachyury* and *Pax6*, respectively) were similarly induced in *Klf5-overexpressing* and control EBs tested. Data were generated from 2 independent, passage-controlled ESC lines. Error bars = s.d.; *Cdx2* (day 6), \**P* = 0.0496, df = 2, t = 4.321; *Gata4* (day 9), \**P* = 0.0423, df = 2, t = 4.704. **E.** Single GFP-labeled, *Klf5-overexpressing Dux^-/-^* ESCs exhibit a bi-potential cell fate in chimeric blastocysts. ESC contribution to the ICM and/or the TE was determined by the localization of GFP-positive ESC progenies (left). The percentage of chimeric blastocysts with ESC contribution to the ICM, the TE, or both were quantified (right). Scale bar, 20 μm. **F, G**. Human DUX4 directly induces *KLF5*, but not other 2C-transcription factors (*DPPA2* and *NELFA*). **F**. Re-analysis of RNA-seq data from *DUX4*-overexpressing and control ESCs suggests that the DUX/KLF5 regulation, but not the DUX/DPPA2 or DUX/NELFA regulation, is conserved in hESCs, *KLF5*, \*\*\*\**P* < 0.0001. **G.** Read density plots for DUX4 ChIP-seq data in hESCs(Hendrickson et al., 2017) demonstrate its enriched occupancy proximal to *KLF5*, but not *DPPA2* or *NELFA*. **H.** Our proposed model for the role of Klf5 in promoting bi-potential cell fate in preimplantation embryos. 2C-specific transcription factors, such as Dux, are induced in the early 2C stage, and converge on transcriptional activation of *Klf5.* Klf5 establishes bi-potential cell fate in *vitro* and in *vivo* through its dual regulation of ICM (in cooperation with *Klf4*) and TE specification genes.

In most bi-potential ESCs with expanded cell fate potential, an aberrant increase in *Dux* constitutes the key mechanism underlying the MERVL induction (Choi et al., 2017; Hu et al., 2020; Macfarlan et al., 2012; Yan et al., 2019). In contrast, Klf5 induced MERVL induction and bi-potential cell fate in ESCs likely acts downstream of *Dux*. *Klf5* overexpression invariably failed to induce *Dux* (data not shown); MERVL induction by *Klf5* was preserved even in a *Dux* knockout background (Fig. 4C). More importantly, EBs derived from *Klf5-overexpressing Dux^-/-^* ESCs, but not control *Dux^-/-^* ESCs, induced markers for both extraembryonic lineages (TE marker *Cdx2* and PrE marker *Gata4*) and embryonic lineages (Ectoderm marker *Pax6,* Mesoderm marker *Brachyury* and Endoderm marker *Foxa2*) (Fig 4D, Sup. Fig. S4C). And a single *Klf5-overexpressing Dux^-/-^* ESC was able to colonize the ICM, TE or both in 44%, 22% and 33% chimeric blastocysts, respectively (Fig. 4E).

The functional importance of Klf5 and its regulation by Dux prompted us to investigate the evolutionary conservation of the Dux/Klf5 axis between mouse and human. Mouse *Dux* and human *DUX4* are both induced at the onset of ZGA to govern the induction of early zygotic genes (Hendrickson et al., 2017; Iaco et al., 2017). We then investigated to what extent Dux regulation on *Klf5* was conserved between mouse and human. We analyzed published RNA-seq and ChIP-seq data for *DUX4* over-expressing human ESCs (hESCs) (Hendrickson et al., 2017). Intriguingly, human *KLF5* was significantly upregulated by *DUX4* in RNA-seq analysis, yet a subset of bi-potency regulators and known Dux targets in mouse, such as *NELFA* and *DPPA2*, were not impacted by enforced *DUX4* expression (Fig. 4F). This is consistent with ChIP-seq data, as DUX4 demonstrated enriched occupancy in multiple regulatory regions proximal to the *KLF5* transcriptional start site, whereas no DUX4 occupancy could be detected near *NELFA* and *DPPA2* (Fig 4G). The evolutionary conservation of the Dux/Klf5 axis, but not Dux/Dppa2 or Dux/Nelfa axis, highlights the functional importance of Klf5 in establishing a bi-potential cell fate in early blastomeres.

While Klf transcription factors are essential for bi-potential cell fate, many 2C specific transcription factors likely act redundantly, and the individual disruption of *Dux*, *Dppa2*/*4*, *Tbx3* and *Nelfa* in knockout mice failed to yield a preimplantation phenotype ((IMPC), 2014; Chen and Zhang, 2019; Nakamura et al., 2011). This contrasts to the strong ICM and TE specification defects when *Klf5* and *Klf4* are both knocked down (Fig. 3I, 3J). Hence, it is likely that multiple 2C specific transcription factors converge on Klf5 via evolutionarily conserved mechanisms, which promotes major 2C-specific ERV expression and establish bi-potential cell fate (Fig. 4H).

## Discussion

Bi-potential blastomeres exhibit an intrinsic transcriptional program co-expressing both ICM and TE genes at a modest level; subsequent extrinsic signaling then acts as the decisive factor to promote one lineage while repressing the other (Pfeffer, 2018). Hence, the molecular nature of the bi-potential plasticity is likely a cell state with the developmental plasticity to activate either ICM or TE specification genes, in a context-dependent manner. To elucidate the key molecular pathway regulating the bi-potent cell fate, we identified 3 major ERV families as the molecular hallmarks for bi-potential blastomeres. The LTR sequences of all three 2C-ERVs likely share critical regulatory sequences for key bi-potential transcription factor(s). We performed motif analyses on these LTRs and identified Klf5 as an important regulator for 2C-specific ERVs and 2C-like bi-potential cell fate.

Upon overexpression, Klf5 establishes a robust bi-potential cell fate in single ESCs to yield multiple terminally differentiated embryonic and extra-embryonic lineages in chimeric mid-gestation embryos. This effect of *Klf5* is highly dependent on its expression threshold, as the endogenous *Klf5* expression in pluripotent ESCs is not sufficient to confer bi-potential cell fate, yet an elevated expression Klf5 level is. The ability of Klf5 to establish a bi-potential cell fate in ESCs is consistent with its functional importance in preimplantation cell fate decisions. *Klf5* is expressed in both ICM and TE, with a strong enrichment in TE. *Klf5* deficiency alone impairs TE specification, and *Klf5* and *Klf4* deficiency in combination impairs both TE and ICM specification (Fig. 4H).

While functional redundancy exists among preimplantation-specific Klf transcription factors, Klf5 is the most important Klf for bi-potential cell fate. Klf5 is the only known Klf factor expressed in both ICM and TE, with a knockout phenotype impairing blastocyst cell fate specification (Azami et al., 2017; M. et al., 2008). Klf5 alone is necessary and sufficient to promote TE specification, whereas Klf5 acts redundantly with other Klf factors, such as Klf4, to promote ICM specification in preimplantation embryos. Hence, Klf5 acts at the core of a robust Klf transcription network, which promotes dual induction of ICM and TE specification genes to establish bi-potential developmental plasticity in cleavage stage blastomeres.

Findings from our *Klf5* loss-of-function studies are seemingly different from the conclusion of a previous study, which reported defects in both ICM and TE cell fate decisions in *Klf5* knockout embryos (Ema et al., 2008). Nevertheless, the actual data in this study, as well as those from a later study(Lin et al., 2010), clearly indicated that *Klf5* knockout caused a strong and fully penetrant TE defect, but a mild, lowly penetrant ICM defect. The small difference in the extent of ICM defects in different *Klf5* loss-of-function studies can be attributed to the different genetic background. These observations are intriguing, as Klf5 induces both ICM and TE genes, yet its deficiency preferentially impacts the TE cell fate. As is clear from our studies, Klf5 acts alone to promote the TE cell fate, and Klf5 functions redundantly with Klf4 to regulate ICM cell fate.

Klf5 induction is regulated by multiple transcription factors that regulate 2C-specific transcriptome, including Dux, Dppa2 and Tbx3. Unlike *Klf5*, none of these 2C transcription factors exhibit strong preimplantation phenotype in knockout studies((IMPC), 2014; Chen and Zhang, 2019; Nakamura et al., 2011). It is likely that they act redundantly in promoting bi-potential cell fate, and that their collective regulation on developmental plasticity converges on Klf5, and possibly other Klf genes. In particular, Dux regulation on Klf5 is evolutionarily conserved between mouse and human, but a number of well characterized, 2C-specific Dux targets in mouse are not regulated by human DUX. Our findings suggest an evolutionarily conserved functional importance of the *Dux-Klf5* axis in regulating the developmental plasticity of early blastomeres in mammals.

Bi-potential cell fate is a complex biological state, which may not be always associated with 2C-ERV induction. Expanded pluripotent stem cells (EPCs), derived either from mouse 8C blastomeres or from treatment of a chemical cocktail, yield embryonic and extra-embryonic potency in chimeric embryos, without inducing MERVL (Yang et al., 2017a, 2017b). Future studies will likely identify additional pathways acting in parallel to Klf5 to promote this developmental plasticity.

## Acknowledgments

We thank Nina Xiong, Jessica Mar, Andrew Modzelewski, Joseph Martin, Pratishtha Rawat, Suifang Mao, Yun Zhou and Sean Chen for technical assistance, discussion, input and critical evaluation of the assumptions and evidence provided in this study. Further, we express our deepest gratitude to Alberto de Laco and the Trono lab for their generosity in sharing *Dux* knockout and control ESC lines. This study utilized Berkeley’s Vincent J. Coates Genomics Sequencing Laboratory at UC Berkeley (supported by NIH S10 OD018174 Instrumentation Grant) and computational resources provided by (Berkeley Statistics, Biostats and UT-dallas). : L.H. is a Thomas and Stacey Siebel Distinguished Chair Professor, a Bakar fellow, who is supported by a Howard Hughes Medical Institute (HHMI) Faculty Scholar award, and several grants from the National Institutes of Health (NIH; 1R21HD088885, GRANT12095758 1R21OD027053-01). M.K. is supported by a CRCC pre-doctoral fellowship.

## Author contributions

M.K. has conceived, designed and executed the majority of the experiments reported in Figure 1, 3, and 4. Y.C. performed most experiments reported in Figure 2. M.K., K.L., H.B. and Z.X. provided bioinformatic analyses on preimplantation single cell RNA-seq analysis, motif enrichment analysis and visualization, ChIP-seq analysis and visualization, and ingenuity pathway enrichment analysis. C.C prepared ChIP-seq samples and M.K. MERVL-reporter ESC data was generated by N.P. Teratoma experiments and analysis were done by Y.J., R.V. and M.K. Morula injections to generate chimeric blastocysts as well as all imaging and quantification were performed by S.Y., Y.C., and M.K. Depletion experiments by RNAi and CRISPR-EZ as well as downstream qPCR and immunofluorescence was done by M.K. Data mining of CistromeDB was done by K.L and M.K.;

## Data and materials availability

The ChIP-seq data generated for this publication have been deposited in NCBI’s Gene Expression Omnibus and are accessible through GEO Series accession number GSE137036 (https://www.ncbi.nlm.nih.gov/geo/query/acc.cgi?acc=GSE137036, reviewers’ token: cfgnoowkzdytdol). All other data is can be found in the paper or supplementary materials.

## Supplementary Materials for

### Materials and Methods

#### RNA-seq

Preimplantation RNA-seq analyses were performed on previously published data (GSE45719)(Deng et al., 2014), specifically the single cell samples from the zygote to blastocyst developmental stages. Bulk RNA-seq analyses were also performed on previously published data (GSE85632)(Hendrickson et al., 2017). Fastq files were processed using Kallisto, and the resulting gene count matrix was generated(Bray et al., 2016). Genes were filtered, keeping only those with at least 300 counts across 9 samples. We then used the limma *R* package to perform full-quantile normalization(Ritchie et al., 2015). Or DESeq2 to test for differential expression(Anders and Huber, 2010). One sample was filtered out based on the PCA plot, leaving 258 samples and 12909 genes. Pseudotime was then computed using the *R* package slingshot (Street et al., 2018). Trends of gene expression were obtained using tradeSeq (Van den Berge et al., 2020).

For quantification of retrotransposons, Fastq files were analyzed by STAR with options ‘--outSAMtype BAM SortedByCoordinate --outSAMattributes XS --outFilterMultimapNmax 100000000’(Dobin et al., 2013). The repeatmasker track was downloaded from the UCSC Table browser, using the GRCm38/mm10 reference genome and filtered for short repeats. The genome annotation was downloaded from Gencode (Basic gene annotation, release M22)(Harrow et al., 2012). The bam files were then compared with the repeatmasker or the basic annotation using FeatureCounts, with options ‘-O -p -B -C -M’(Liao et al., 2014). Using the pseudotime obtained on the as described above, the RT expression of a few selected families across time was then plotted.

#### Motif Enrichment Analyses

All annotated LTR sequences for MERVL, ORR1A0 and ORR1A1 in the C57B/6 mouse genome were downloaded from DFAM(Hubley et al., 2016). Subsequently, HOMER was used to generate scrambled control sequences via the HOMER’s scrambleFasta.pl script. And the HOMER known module was used to compute motif enrichment within the LTRs of the tested ERVs for all known transcription factor motifs with the -opt flag to optimize the degeneracy threshold to get the best enrichment(Benner et al., 2017). The analyses were performed on each 2C-ERV family individually as well as a merged cohort. For visualization of specific motif instances within gene bodies, genomic sequences were obtained from the UCSC mm10 genome browser and scanned with the JASPAR database web client with default parameters(Mathelier et al., 2016).

#### Luciferase Assays

For MERVL-luciferase reporter assays, we used the pGL3 luciferase reporter vectors (Promega, Cat. # E1751) that harbors the MERVL_125-375_-fragment previously described (MERVL-Luc)(Choi et al., 2017). MERVL-Luc reporters and control Renilla luciferase reporter pRL-TK (Promega, Cat. # E2241) were co-transfected into ESCs (600 ng and 150 ng per well of a 12-well plate, respectively), using Lipofectamine 2000 (Life Technologies, Cat. # 11668027). Transfection complexes containing the reporter constructs were prepared in Opti-MEM Reduced-Serum Medium (Life Technologies, Cat. # 31985062). After trypsinization with 0.25% Trypsin + EDTA (Life Technologies, Cat. # 25200-056), 100,000 cells were resuspended in ES media lacking Pen Strep, incubated with transfection complexes for 10 minutes at 37° C, and then transferred to one well of a 12-well plate containing irradiated MEF feeders. After 48 hours, transfected ESCs were trypsinized, plated onto gelatin-coated plates for 1 hour to remove feeders, and then assayed for luciferase activity by Dual-Luciferase® Reporter Assay System (Promega, Cat. # E1910) using a Glomax 20/20 Luminometer (Promega).

#### Derivation and culture of mouse ESCs

Mouse ESCs were isolated based on published protocols with slight modifications(Bryja et al., 2006). Uteri containing E3.5 wild-type embryos were isolated from timed pregnancies, and subsequently put in Knockout DMEM (Life Technologies, Cat. # 10829-018) supplemented with 10mM HEPES (Life Technologies, Cat. # 15630-080). E3.5 blastocysts were flushed with 1ml syringes with 18G needles, and individually transferred to a 12-well plate seeded with irradiated MEF (mouse embryonic fibroblasts) feeders in 1 ml N2B27 medium containing 100 U/ml LIF (EMD Millipore, Cat. # ESG1107), 1 µM PD0325901 (Sigma, Cat. # PZ0162) and 3 µM CHIR99021 (EMD Millipore, Cat. # 361559). After 5 days of incubation, embryo outgrowth was separated from the trophectoderm (TE), picked up by a 10 µl pipette, transferred to 20 µl Accutase (Life Technologies, Cat. # A11105-01) and incubated at 37°C for 20 min to dissociate cells. Dissociated cells were then cultured on irradiated MEF feeder cells with N2B27 medium containing LIF and two inhibitors for one passage. Subsequently, ESCs were passaged with 0.25% Trypsin-EDTA and maintained in regular mouse ES medium.

ESCs were cultured onto irradiated MEF feeder layers in the M15 ESC medium, which contained Knockout DMEM (Invitrogen, catalogue no. 10829-018), 15% ES-grade fetal bovine serum (Invitrogen, Cat #. 16141079), 2 mM L-glutamine (Invitrogen, Cat #. 25030-164), 1×10^−4^ M MEM non-essential amino acids (Invitrogen, Cat #. 11140-076), 1×10^−4^ M 2-mercaptoethanol (Sigma, Cat #. M3148) and 1% 100× penicillin and streptomycin. ESCs were split every two days.

#### ESC transfection

To overexpress *Klf5, Dux,* or *Dppa2* in ESCs, cells were transfected with *PiggyBac* vectors containing an EF1α-driven *Klf5, Dux,* or *Dppa2* expression cassette and an Ubc-puromycin selection marker. Individual *PiggyBac*-plasmid was mixed with the *PiggyBac* transposase plasmid in a 1:1 ratio, and subsequently transfected into ESCs using Lipofectamine 2000 (Life Technologies, Cat. # 12566014) following the manufacturer’s instruction. Cells were selected with 3 μg/ml puromycin for two days on puromycin resistant MEF feeders, and then cultured in puromycin-free M15 ES medium for following analyses. The *PiggyBac*-EF1α -GFP-Ubc-Puro plasmid was used as a negative control.

#### Single-Embryo QPCR

All single-embryo cDNA was prepared according to the Single Cell-to-Ct qRT-PCR kit (Life-Technologies, Cat# 4458236) with slight modifications. Pronuclear, 2-cell, 8-cell and blastocyst stage embryos were isolated and passed through three washes of PBS. Single embryos were then placed into individual PCR tubes and lysed in twice the recommended volume of Lysis/DNAse (20 μL) for 15min at room temperature. Then, 2 μL of Stop Solution was added and incubated for 2 min. At this point, half of the reaction was stored in −80°C conditions as a technical replicate and the remaining sample (11 μL) continued through the original Single Cell-to-Ct protocol. All qRT-PCR reactions were performed using SSO Universal SYBR Green SuperMix, as per manufacturer instructions (Biorad, Cat# 1725275). All QPCR analyses were performed on the StepOnePlus Real Time PCR system (ThermoFisher, Cat# 437660).

#### Real-time PCR

RNA was isolated using Trizol following manufacturer’s instruction (Life Technologies, Cat. # 15596). cDNA was reverse-transcribed using iScript Advanced Reverse-Transcriptase (Bio-Rad, Cat. # 1725037). All real-time qPCR analyses were performed using SYBR FAST qPCR Master Mix (Kapa Biosystems, Cat. # KK4604), following manufacturer’s protocol. Real time PCR analyses on retrotransposons detect their expression at the family level, using primers designed from the corresponding consensus sequences. *Actin* was used as a reference for both mRNA and retrotransposon quantitation in real time PCR analyses. All real time PCR primers used in our studies are listed in Supplementary Table S5.

#### Teratoma generation and histological analyses

1×10^6^ of WT or *Klf5*-overexpressing ESCs were injected into the dorsal flanks of 6-7-week-old immune-deficient NCr-nu/nu female mice (Taconic, Cat# NCRNU). After 4-5 weeks, resulting teratomas were collected by surgical removal, fixed overnight in 10% buffered formalin (Fisher Scientific, Cat. # SF100-4), dehydrated in a graded series of ethanol solutions, embedded in Paraplast X-TRA paraffin (Fisher Scientific, Cat. # 23-021-401), sectioned at 6 µm thickness, and stained with hematoxylin and eosin (H&E) using standard procedures(Choi et al., 2011). These paraffin sections will be subjected to immunohistochemistry

#### Embryoid body (EB) differentiation

For EB differentiation, ESCs were plated in 10cm petri dish (150,000 cells/ml) in ESC M15 medium without LIF and were gently cultured on a rotator after removal of feeder cells. Samples were collected at day 0, 3, 6 and 9 post EB differentiation for real-time PCR analyses and for immunofluorescence staining (see below).

For hanging drop EB formation, ESCs colonies were removed from feeders and dissociated to near single cell suspension using trypsin (Life Technologies, Cat # 25200114). Small clumps of cells (10-20 cells/clump) were then mouth pipetted into 25uL drops of M15 medium without LIF and placed on the underside of the lid to a 6cm petri dish. 2mL of PBS was added to the 6mL petri dish to prevent evaporation of the EB culture drops. EBs were cultured for 72h before subjected to fixation for immunofluorescence staining as described below.

#### Generation of chimeric blastocysts and chimeric embryos from ESCs

Klf5 overexpressing wildtype ESCs, *Klf5*-overexpressing *dux^-/-^* ESCs and the corresponding control cells were all engineered to express GFP from a piggybac vector. To generate chimeric blastocysts, single ESCs was injected into each E2.5 8 cell C57Bl/6N wild-type recipient morulae. Injected embryos were then cultured overnight in KSOM (Millipore, Cat # MR-106-D) to obtain chimeric blastocysts. GFP positive cells were scored in the ICM, TE or both in the chimeric blastocyst based on their morphology and location.

To generate chimeric mid-gestation embryos, we initially injected 10-15 GFP labeled, *Klf5*-overexpressing wildtype ESCs into C57Bl/6N wildtype recipient blastocysts, followed by a uterus transfer into the CD1 pseudo-pregnant mothers. Chimeric embryos were then collected at E12.5 for immunofluorescence analyses (see below). Subsequently, we generated E12.5 chimeric embryos by injecting single, GFP labeled ESCs into each recipient blastocysts for the same analyses.

#### Immunohistochemistry and immunofluorescence analyses

For immunohistochemistry (IHC) analyses on teratomas, 6µm paraffin sections were deparaffinized, dehydrated, and subjected to heat-induced antigen retrieval in a pressure cooker using Target Retrieval solution (DAKO, Cat. # S1699). Slides were incubated for 10 minutes with 3% H2O2, blocked for 3 hours with PBS containing 5% BSA and 0.3% Triton X-100, and incubated with primary antibodies against PL-1 (1:75, Santa Cruz Biotechnology, Cat. # sc-34713) overnight in PBS buffer containing 1% BSA and 0.3% Triton X-100. Slides were then incubated with horseradish peroxidase (HRP)-conjugated secondary antibodies for 2 hours at room temperature, and then subjected to 3,3’-Diaminobenzidine (DAB) staining (Life Technologies, Cat. # 00-2014) followed by a counterstain with Mayer’s hematoxylin (Electron Microscopy Sciences, Cat. # 26503-04). The sinusoidal trophoblast giant cells (s-TGCs) were identified by their enlarged nuclei and adjacent location in the maternal blood sinusoid space.

For immunofluorescence (IF), ESC colonies, differentiated EBs and blastocysts were fixed with 4% paraformaldehyde (Electron Microscopy Sciences, Cat # 19202) for 10 min at room temperature and incubated with blocking solution (0.1%Triton X-100 and 5% normal goat serum in PBS) for 1 hour at room temperature. EBs or ESCs were incubated overnight at 4°C with appropriate antibodies, including MERVL-Gag, 1:100, Epigentek, Cat # A-2801-100, Oct4, 1:100, Santa Cruz Biotechnology, Cat. # sc-5279, Cdx2, 1:100, Abcam, Cat. # ab76541, Nanog, 1:100, Cosmo Bio, Cat # REC-RCAB0002PF, Klf5, 1:100, Protein tech, Cat # 21017-1-AP). Subsequently, samples were stained with goat anti-rabbit IgG (H+L) secondary antibody, Alexa Fluor 594-conjugated secondary antibody (1:500, Life Technologies, Cat. # A11037) for 1 hour at room temperature (ESCs) or overnight at 4°C (EBs). Samples were then stained with DAPI (300 nM, Sigma, Cat. # D9564) and subjected to imaging analyses using spinning disk confocal microscopy (Andor CSU-X on Nikon Eclipse TE200-E). ImageJ was used to analyze mean fluorescence intensity of acquired images.

For IF staining of E12 chimeric mouse embryos, samples were fixed with 4% paraformaldehyde (Electron Microscopy Sciences, 19202) for 2 hours, incubated in 30% sucrose (Fisher, Cat # S5-500) overnight at 4°C, embedded in Tissue-Tek O.C.T. compound (VWR, Cat. #25608-930), and cryo-sectioned at 8 µm. These sections were subsequently subjected to IF analyses using antibodies against GFP (1:100, Abcam, Cat. # ab38689), Tpbpa (1:200, Abcam, Cat. # ab104401), and Mtp1 (1:150, Alpha Diagnostic, Cat. # MTP11-A). Trophoblast giant cells were identified based on their unique location in the placenta and their distinct morphology of enlarged nuclei. Spongiotrophoblasts were identified based on the staining of Tpbpa; syncytiotrophoblasts were identified based on the staining of Mtp1. The bilaminar structure of the yolk sac is identified by DAPI staining, and the visceral endoderm is identified by its columnar, epithelial morphology.

#### Chromatin immunoprecipitation (ChIP) and ChIP-seq libraries

V5 ChIP assays were performed using ESCs overexpressing a V5-tagged Klf5 protein. Wild-type D3 mouse ESCs were used as a control. Cells were cross-linked for 5’ at room temperature with 1% formaldehyde-containing Knockout D-MEM; cross-linking was stopped by PBS-glycine at a final concentration of 0.125 M. Cells were washed twice with ice-cold PBS, scraped, centrifuged for 10 min at 4000 rpm and flash-frozen in liquid nitrogen. Cell pellets were thawed in ice, resuspended in cell lysis buffer (5 mM PIPES, pH 8.0, 85 mM KCl, and 0.5% NP-40, 750 μl/15 cm plate) and incubated for 10 min on ice. During the incubation, the lysates were repeatedly pipetted up and down every 5 minutes. Lysates were then centrifuged for 10 min at 4000 rpm. Nuclear pellets were measured and resuspended in 6 volumes of sonication buffer (50 mM Tris-HCl, pH 8.1, 10 mM EDTA pH 8.0, 0.1% SDS), incubated on ice for 10 min, and sonicated to obtain DNA fragments below 2000 bp in length (Covaris S220 sonicator, 20% Duty factor, 200 cycles/burst, 150 peak incident power, 32 cycles of 20” on and 40” off). Sonicated lysates were cleared by centrifugation (20’ at 13200 rpm) and 800 μg of chromatin were diluted in RIPA buffer (10 mM Tris-HCl, pH 8.0, 1 mM EDTA pH 8.0, 0.5 mM EGTA, 1% Triton X-100, 0.1% SDS, 0.1% Na-deoxycholate, 140 mM NaCl), precleared with Protein G sepharose (GE Healthcare, Cat # GE17-0618-01) for 2 hours at 4°C and immunoprecipitated overnight with 8 μg of normal mouse IgGs (ChromPure mouse normal IgG; Jackson ImmunoResearch), or anti-V5 antibodies (Invitrogen R960-25). 4% of the precleared chromatin was saved as input. After the overnight incubation, samples were incubated with 25 μl Protein G sepharose beads precleared overnight in RIPA buffer with 0.5% (w/v) BSA and incubated for 2 hours at 4°C. Immunopecipitated samples were washed 5 times with RIPA buffer, once with LiCl buffer (0.5% NP-40, 0.5% Na-deoxicholate, 250 mM LiCl, 1 mM EDTA pH 8.0), and once with TE. After the last wash, immunoprecipitated complexes were eluted from the beads twice with 150 μl of TE with 1% SDS, each time incubating 30 min in a thermomixer set at 37°C and 900 rpm. The 300 μl eluted material was added of 1 μl RNaseA (10 mg/ml) and 18 μl 5M NaCl and incubated at 67°C for 4-5 hours to reverse formaldehyde cross-linking. Inputs were added of elution buffer to 300 μl total volume, and subject to the same treatment. Reverse cross-linked samples were added of 2.5 volumes of ice-cold ethanol and precipitated overnight at −20°C. DNA was pelleted by centrifugation (20min at 13,200 rpm and 4°C), and pellets resuspended in 100 μl TE, 25 μl 5X PK buffer (50 mM Tris-HCl, pH 7.5, 25 mM EDTA pH 8.0, 1.25% SDS), and 1.5 μl proteinase K (20 mg/ml), and incubated 2 hours at 45°C. After proteinase K digestion, DNA was purified with the Qiagen QIAquick PCR Purification Kit, eluted in 60 μl of water and analyzed by qPCR together with 2% of the input chromatin prior to ChIP-seq library preparation using SYBR FAST qPCR Master Mix (KAPA, Biosystems Cat. # KK4604).

ChIP-seq libraries were prepared using the Illumina TruSeq™ DNA sample preparation kit according to manufacturer instructions with few modifications. We used 150 ng of ChIP input DNA (as measured by Nanodrop) and 50 μl of immunoprecipitated DNA as a starting material; library samples were enriched through 12 cycles of PCR amplification. We assessed library quality and fragment size by qPCR and Fragment analyzer™ and sequenced the multiplexed libraries on one lane the Illumina HiSeq4000 sequencing platform (single end-reads, 50 bp long) at the Vincent J. Coates Genomics Sequencing Laboratory at UC Berkeley.

#### ChIP-seq analyses

Fastq files were aligned using STAR with the same options as above. Bam files for Klf5 ChIP samples were merged separately. Peaks were then called using macs2 (Zhang et al., 2008).

Coverage of the peaks by the initial bam files was then computed using betools_2.25.0. Finally, peaks were annotated using the HOMER_4.10 annotatePeaks function, with option -m klf5.motif to specifically look for peaks containing the Klf5 motif. RNA-Seq data was processed as indicated for the TFs plots. For each peak, we therefore also looked at the mean gene-expression (TPM) between late 2-cell and 16 cells (with log1p transformation afterwards) of the gene that is nearest to the peak. Published data were analyzed from GSE85632 (Dux, DUX4), GSM515664 (Nelfa), GSE60066 (Tbx3), GSE117171 (Dppa2).

#### Cistrome DB analyses

Peak bed files from all ChIP-seq experiments performed in mouse ESCs were downloaded from the CistromDB batch repository and subsequently annotated with HOMER2’s annotatePeaks.pl script(Mei et al., 2017). Following annotation, factors with the potential to regulate Klf5 were defined as those that had peaks present within a window 2.5kb upstream and 2.5kb downstream of the TSS.

#### *Klf5* knockdown in preimplantation embryos using RNA interference (RNAi)

5IU of PMSG (Prospec Cat# HOR-272) and HCG (Sigma-Aldrich, Cat # CG5-1VL) were injected intraperitoneally into 4-5 week old, wild-type C57BL/6J female or or C57BL/6J C3H/HeJ F1 mice 46-48h apart. Immediately following HCG injection, females were paired with C57BL/6J stud males and pronuclear stage embryos together with cumulus cell clusters were harvested from plugged females 20h after the mating. Pronuclear stage embryos were dissociated from cumulus cells using Hylorondase (Fisher, Cat # MR-0510F). Embryos were then incubated at 37C with 5% CO2 while siRNAs were prepared for injection. Before injection, scrambled siRNA (Thermo Fisher Scientific, Cat# AM4611) and *Klf5* siRNAs (Thermo Fisher Scientific, Cat# 160900) and *Klf4* siRNAs (Thermo Fisher Scientific, Cat# 156021) were prepared at a working concentration of 100uM were spun at 10k rpm for 5 min to clear debris. In experiments when we double knocked down *Klf4* and *Klf5*, we prepared siRNA mixtures containing 50 uM Klf4 siRNAs and 50 uM Klf5 siRNAs. After the microinjection of siRNAs, embryos were cultured in vitro to E4.0, and then subjected to IF analyses described above.

#### *Klf5* knockout in preimplantation embryos using CRISPR-EZ

C57BL/6J or C57BL/6J C3H/HeJ F1 mice were crossed to C57BL/6J stud males and pronuclear stage embryos were dissociated from cumulus cells using Hylorondase (Fisher, Cat # MR-0510F). After the zona was weakened with acid Tyrode’s (Sigma Aldrich, Cat # T1788), the embryos were subsequently washed in M2 buffer. Cas9 RNP complexes were then assembled *in vitro* by combining 40uM of Cas9 protein with 2ug of sgRNAs targeting Klf5 (CCAGACCGUCCAUGCCCACG, AGCACCCGCGUGGGCAUGGA, GGUCAGCACCCGCGUGGGCA, Synthego). Assembled RNPs were then mixed with a cohort of 50-75 embryos, and electroporated with standard parameters(Modzelewski et al., 2018). Electroporated embryos were then cultured in KSOM until E3.5, at which point they were processed for IF analysis described above.

## Supplementary Table Legend

**Supplementary Table S1. Known motif analysis performed using all ORR1A0, ORR1A1 and MT2_Mm LTR sequences compared against scrambled control sequences.** HOMER motif analysis results is performed for MERVL-LTR (MT2_MM), ORR1A0 LTR and the ORR1A1 LTR. Analyses on separate families as well as the cohort together are shown.

**Supplementary Table S2. Phenotypical analyses of chimeric embryos generated from wild-type and *Klf5*-overexpressing ESCs.**

**Supplementary Table S3. Annotated peaks from Klf5 ChIP-seq experiments using Klf5-overexpressing ESCs.** A list of Klf5 bound regions are shown in BED format. The coordinates are annotated with exact feature information, nearest gene transcriptional start site and the number of Klf5 motifs within the peak.

**Supplementary Table S4. A summary of transcription factors with proximal binding to cis-regulatory sequences of *Klf5***

**Supplementary Table S5. Sequences of real-time PCR primers and sgRNAs.**

## Supplementary figure legend

**Supplementary Figure S1.**
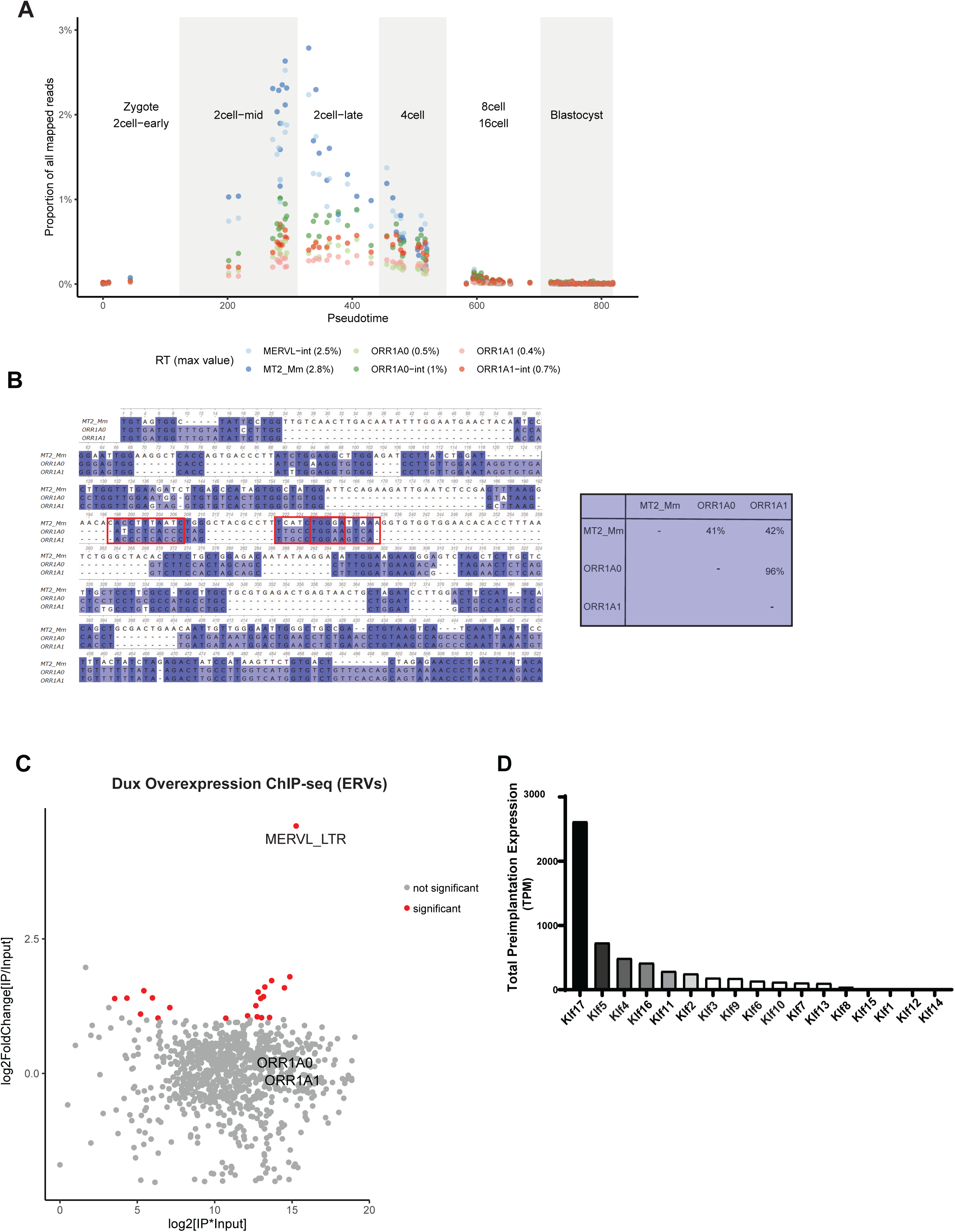
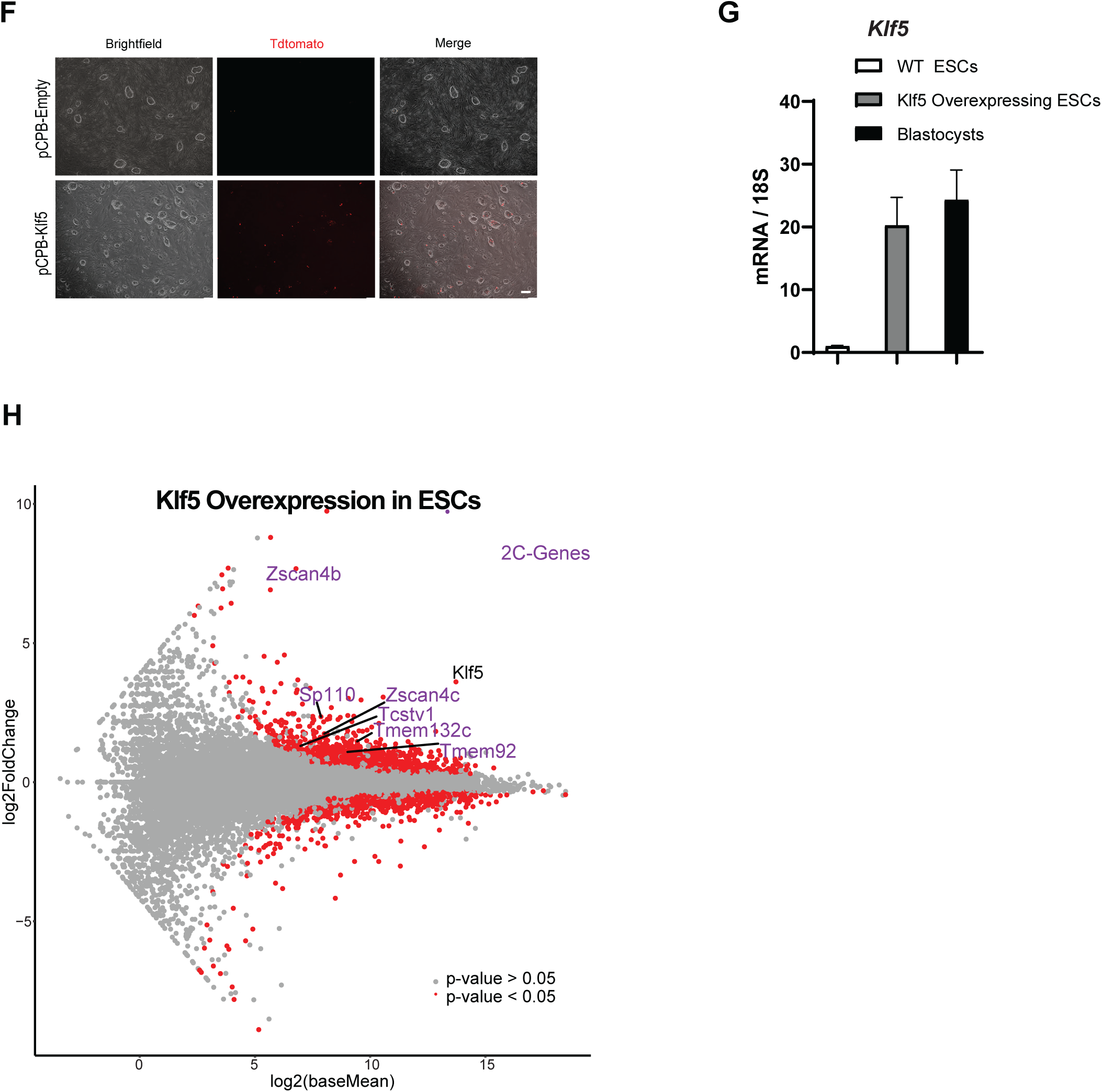
Klf5 directly regulates three major 2C-specific ERV families. **A.** QPCR comparison of *Klf5* expression in WT ESCs, *Klf5* overexpressing ESCs and WT blastocyst embryos demonstrating that the level of *Klf5* overexpression is within a relevant physiological range. **B**. A Pseudotime plot showing the percentage of all reads that mapped to each of MERVL, ORR1A0 and ORR1A1 in preimplantation development. RNA-seq data from Deng et al. are used for this analysis. **C.** Multiple sequence alignment of MERVL, ORR1A0 and ORR1A1 consensus LTR sequences showing the sequence similarity among the three ERV families. ORR1A0 and ORR1A1 are highly homologous in sequences, while MERVL contains stretches of shared sequences with ORR1A0 and ORR1A1. **D.** Diagrams illustrating Klf5 ChIP occupancy at 2C-ERVs that act as alternative promoters for 2C-gene expression. **E.** Dux specifically binds to MERVL, but not ORR1A0 or ORR1A1. An MA-plot was shown to indicate the retrotransposon families enriched for Dux binding using ChIP-seq data of Dux overexpression ESCs from Hendrickson et al. **F.** A bar plot ranking Klf family members by their total expression level (TPM) across all preimplantation developmental stages using published RNA-seq dataset(Deng et al., 2014). *Klf17*, *Klf4*, and *Klf5* are the three most highly and dynamically expressed Klf transcription factors in mouse preimplantation embryos. **G**. *Klf5* overexpression induces MERVL expression in a subset of 2C::Tdtomato wildtype ESCs. Representative images were shown for cultured 2C::Tdtomato ESCs transfected with control and *Klf5* expressing piggyback vectors. **H.** MA-plot of RNA-seq data in control versus *Klf5* overexpressing ESCs showing the upregulation of several well-recognized 2C genes.

**Supplementary Figure S2.**
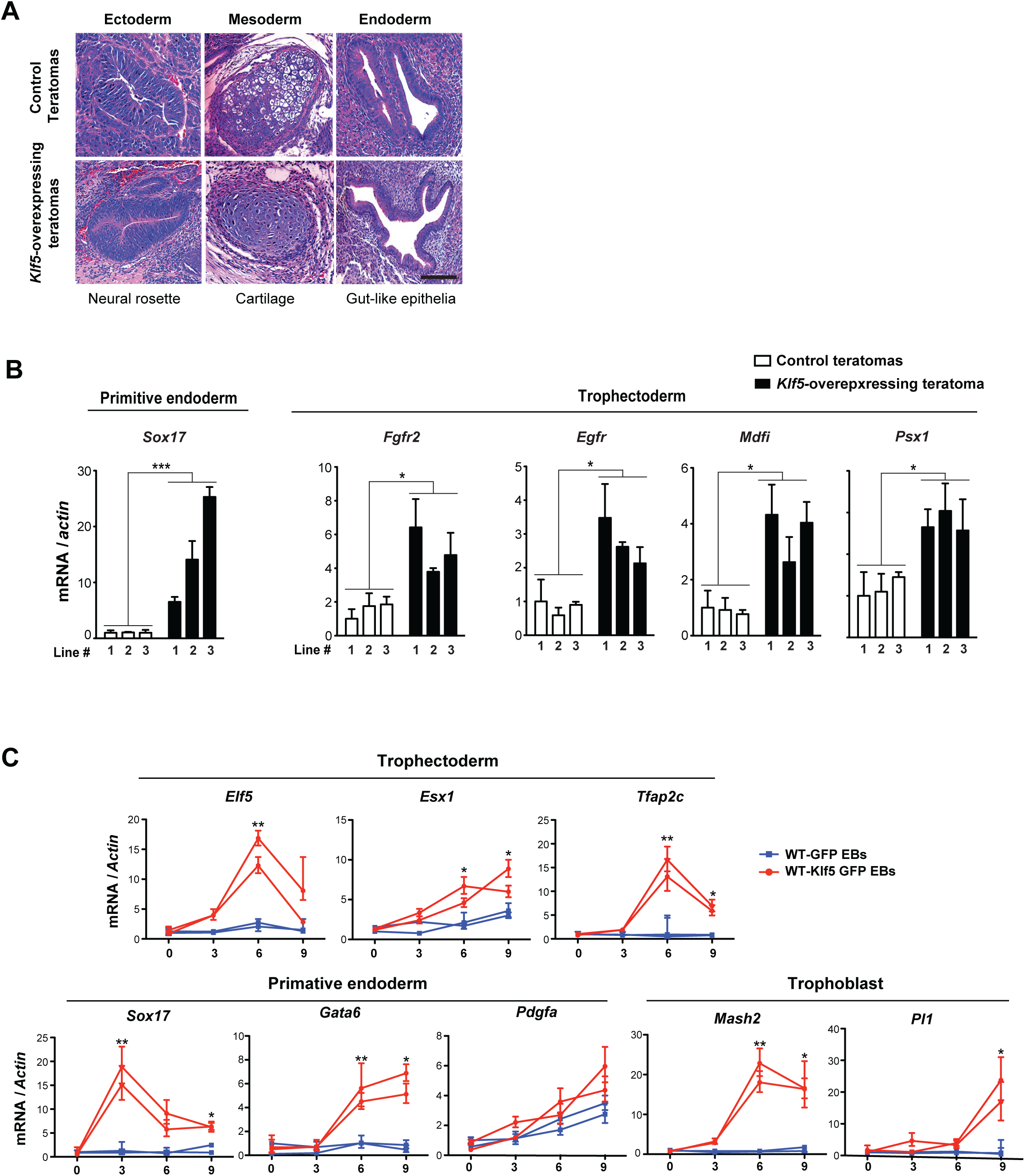
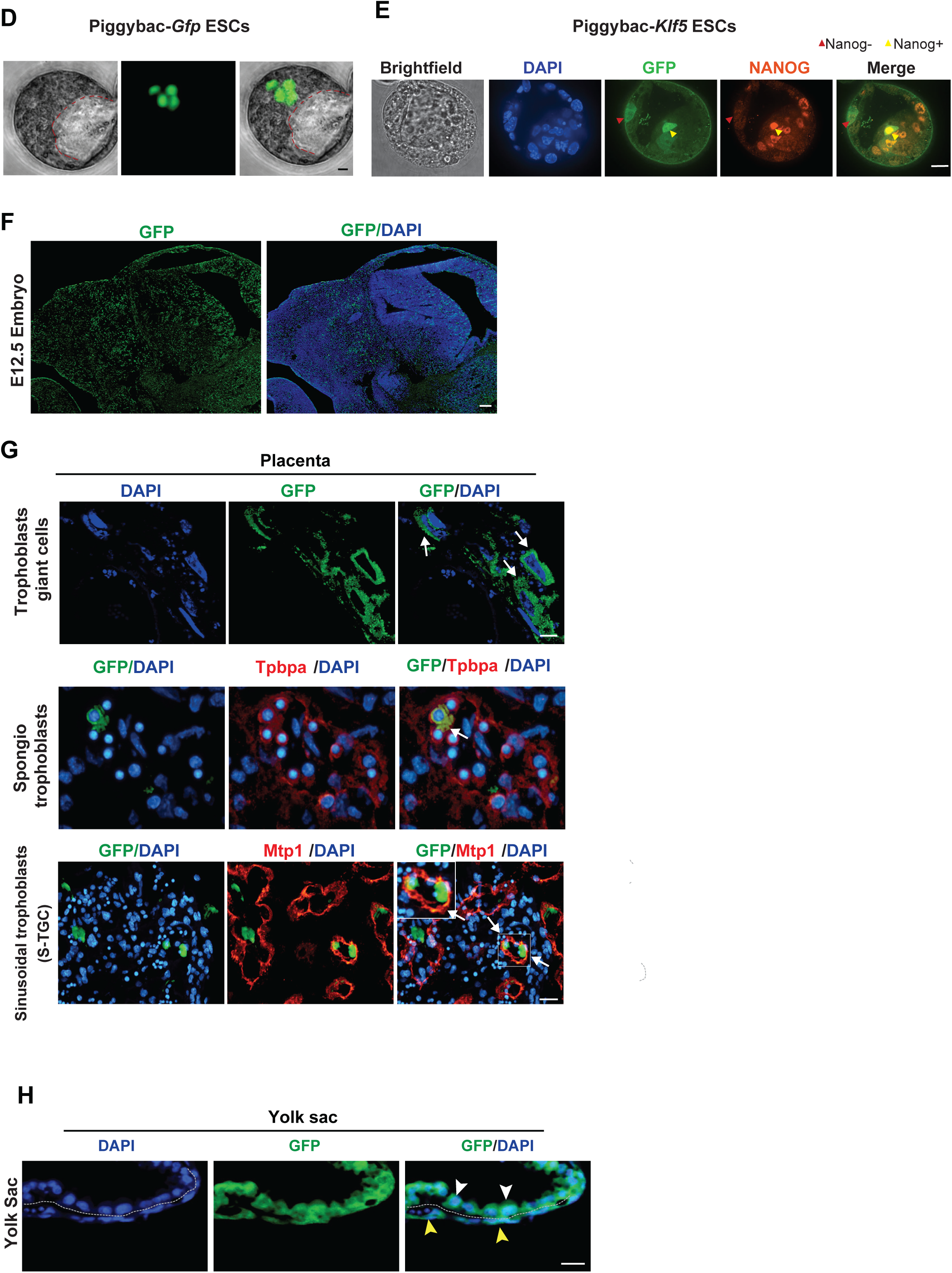
Klf5 confers a bi-potential cell fate in ESCs. **A.** Control and *Klf5*-overexpressing ESCs generate differentiated teratomas containing tissues representative of the three embryonic germ layers (ectoderm, mesoderm and endoderm) as shown by H&E staining. Scale bars, 50 µm. **B.** A real time PCR panel of primitive endoderm (PE) marker (*Sox17*) and TE markers (*Fgfr2, Egfr, Mdfi* and *Psx1*) confirms that Klf5-overexpressing ESCs, but not control ESCs, are capable of robustly activating extra-embryonic transcriptional trajectories in differentiated teratomas. Error bars, *s.d.*, * *P* < 0.05, *** *P* < 0.001. **A**, **B**. A total of three independent pairs of passage-controlled ESCs were compared in the teratoma assay. **C.** A real time PCR panel performed during EB differentiation showed that *Klf5*-overexpressing EBs, but not control EBs, differentiate towards PE (*Sox17),* TE (*Elf5, Esx1* and *Tfap2c*) and more mature trophoblast-like lineages (*Mash2* and *Pl1)*. * *P* < 0.05, *** *P* < 0.001. All *P*-values were calculated using unpaired, two-tailed Student’s *t*-test. **D.** Representative images of a chimeric blastocyst generated by microinjecting a single Piggybac-*Gfp* transduced wildtype ESC into a recipient morula. The progenies of the GFP-labeled ESC only contributed to the ICM. Scale bar = 20µm, red dashed line indicates blastocoel cavity. **E.** The progenies of GFP-labeled *Klf5* overexpressing ESCs can localize to both the ICM and TE compartments. Progeny that localize within the TE compartment lose NANOG protein expression, consistent with a shift from ICM to TE specification whereas progeny in the ICM regain NANOG expression. Red arrows indicate NANOG-progeny in the TE, yellow arrows indicate NANOG+ progeny in the ICM. Scale bar = 20um **F, G.** Klf5 overexpression in ESCs confers a bi-potential cell fate in E12.5 chimeric embryos. **F.** Progenies from Klf5-overexpressing ESCs contribute to embryonic cell lineages in E12.5 chimeric embryos. E12.5 chimeric embryos were generated by injecting either a single ESC or 10-15 ESCs into the recipient blastocysts. Scale bar = 500 µm. **G.** 10-15 GFP-labeled Klf5-overexpressing ESCs were microinjected into each 8 cell morula, followed by embryo transfer into pseudo-pregnant mothers to generate E12.5 chimeric embryos. GFP positive progenies from Klf5-overexpressing ESCs generated trophoblast giant cells (white arrows, identified by their unique cell morphology), spongiotrophoblasts (yellow arrows, identified based on their co-expression of Tpbpa), and syncytiotrophoblasts (orange arrows, identified based on their co-expression of Mtp1). Scale bar = 100µm. **H.** GFP-labeled Klf5-overexpressing ESCs yield both embryonic and extra-embryonic yolk sac lineages. White arrowheads indicate visceral endoderm cells identified based on the bilaminar structure of the yolk sac and their characteristic columnar epithelial morphology. Yellow arrowheads indicate mesothelium derived from embryonic mesoderm. Scale bar, 20 µm.

**Supplementary Figure S3.**
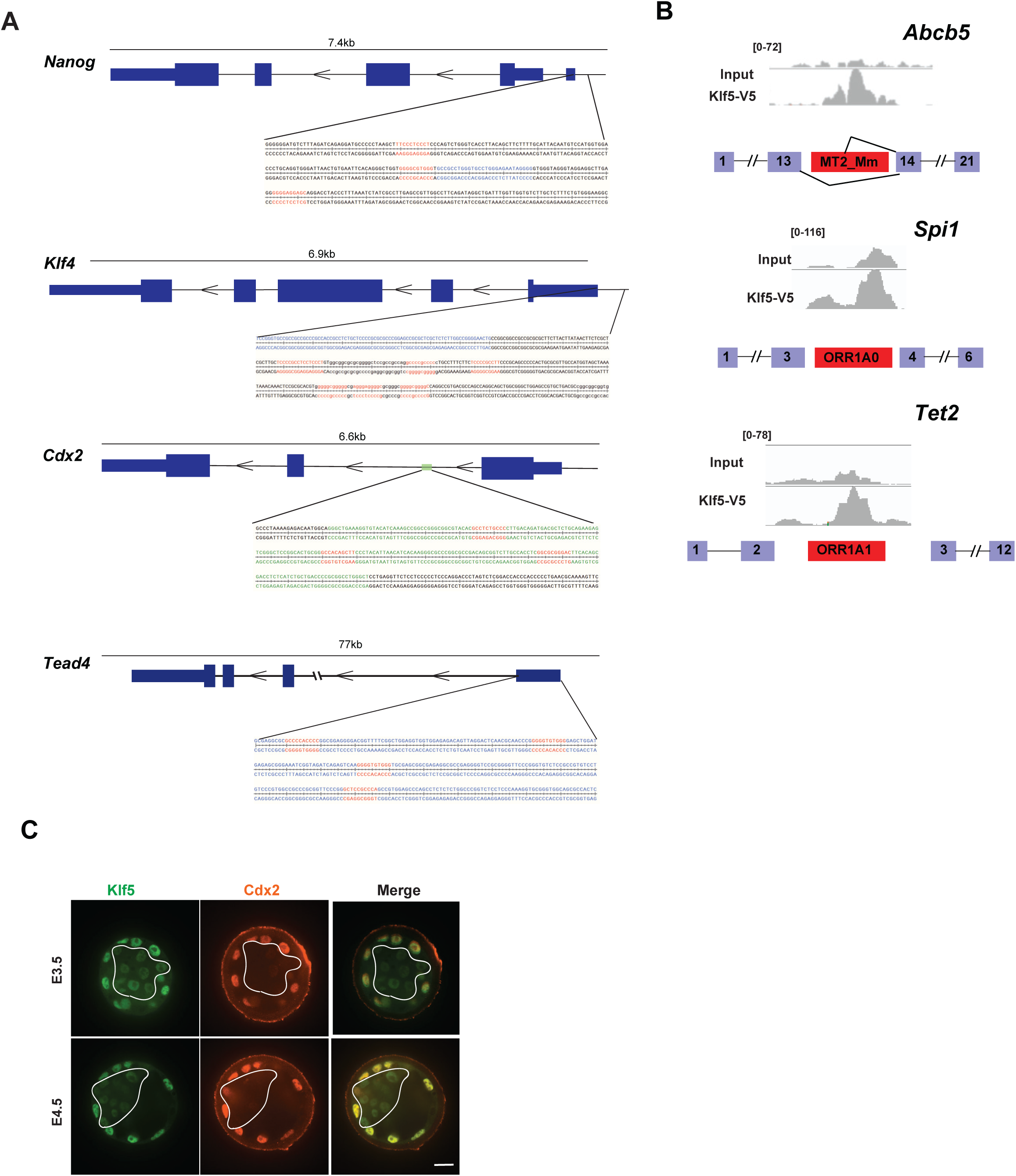
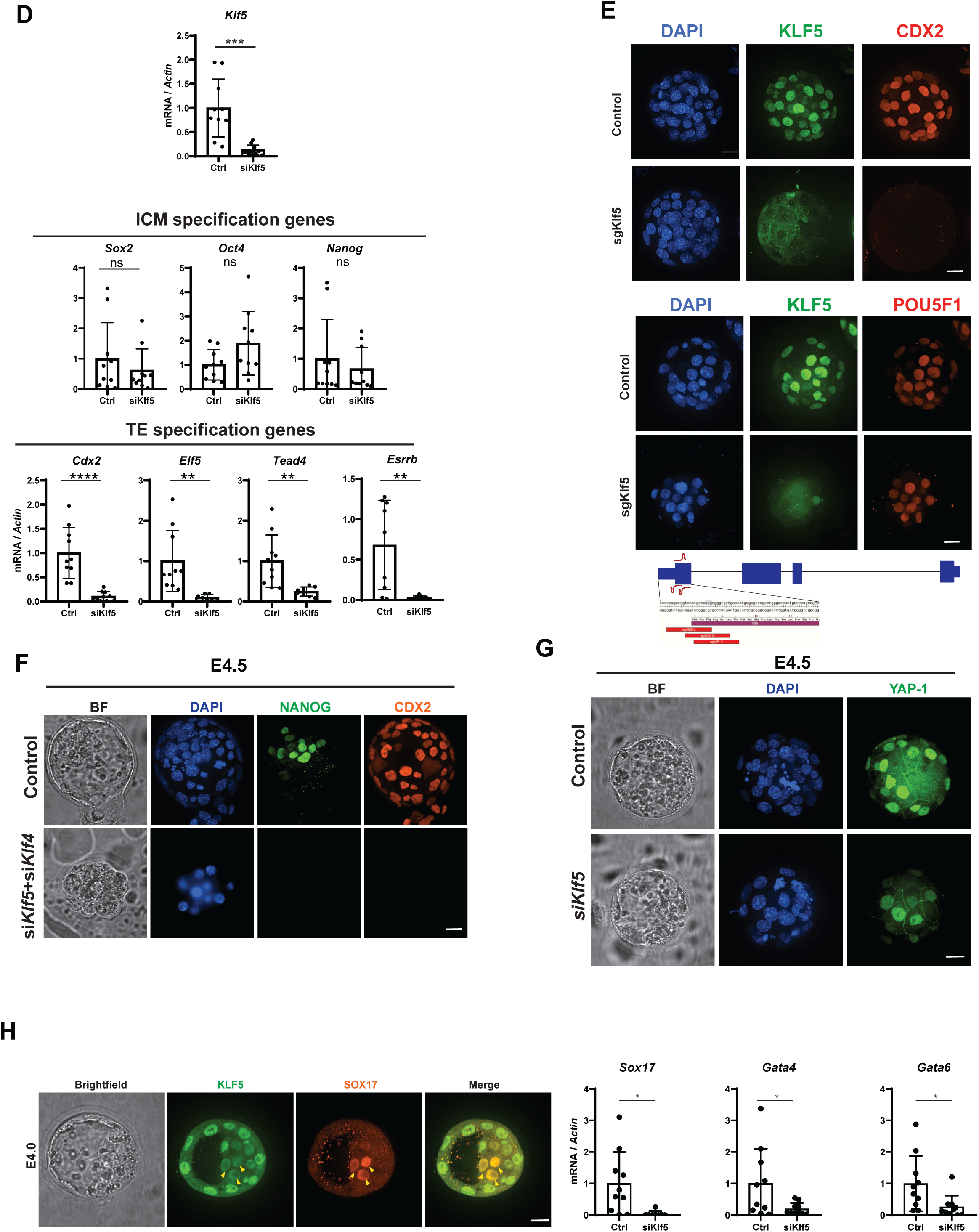
Klf5 acts at the branch point between embryonic and extra-embryonic commitment. **A.** Cis-regulatory elements of *Oct4, Nanog*, *Cdx2* and *Tead4* contain multiple predicted Klf5 binding sites within ChIP-seq peaks identified from *Klf5*-overexpressing ESCs. Blue, a cis-regulatory region containing the TSS; green, known enhancer region of *Cdx2*(Wang and Shashikant, 2007), *red*, predicted Klf binding sites. **B.** Real time PCR analyses from *siKlf5* or scramble siRNAs treated E4.0 embryos demonstrated the efficiency of *Klf5* knockdown. Error bar = s.d.. *** *P* = 0.0003, df = 18, t = 4.510. The *P*-value is calculated based on unpaired, two-tailed student’s unpaired t-test. **C, D.** *Klf5* knockout cells in blastocysts exhibit a decreased expression of Cdx2 yet retain intact Oct4 expression. Using CRISPR-EZ, we generated E4.0 blastocysts that contained a large subset of *Klf5* deficient cells. Immunofluorescence staining of Klf5 confirming the efficient ablation of *Klf5* by CRISPR-EZ in blastocysts. A marked decrease of Cdx2 in TE was observed in *Klf5* deficient cells in CRISPR edited blastocysts (**C**) yet Oct4 expression remained largely intact (**D**). Scale bars, 20 µm. **E.** Immunofluorescence panel of *Klf5+Klf4* double knockdown embryos showing that downregulation of both factors leads to loss of both NANOG and CDX2 protein in E4.5 embryos prior to apoptosis. **F.** Immunofluorescence panel of YAP-1 in control and *Klf5* knockdown embryos at E4.5 showing both nuclear and cytoplasmic localization of YAP-1 even in si*Klf5* treated embryos, indicated in-tact Hippo signaling despite the loss of *Klf5.* Scale bar = 20um. **G.** Real time PCR analyses confirmed the downregulation of TE specification genes *in Klf5* knockdown blastocyst embryos. Error bars = s.d.; *Cdx2*, \*\*\*\**P* < 0.0001, df = 18, t = 5.297; *Elf5*, \*\**P* = 0.0071, df = 15, t = 3.114; *Tead4*, \*\**P* = 0.0050, df = 16, t = 3.248; *Esrrb*, \*\**P* = 0.0016, df = 17, t = 3.737. * *P* < 0.05, ** *P* < 0.01, *** *P* < 0.001, **** *P* < 0.0001, *n.s*., not significant. All P values were calculated using unpaired, two-tailed student’s t-test. **H.** Comparison of *Klf5* and *Klf4* expression from published mouse preimplantation data (GSE45719) demonstrating that *Klf5* is enriched in the TE and expressed at comparable levels to *Klf4* in the ICM. **I.** Immunofluorescence panel of KLF5 and CDX2 protein confirming transcriptional results that KLF5 is enriched in the TE. White lines show the boundary between the TE and the ICM. Scale bar = 20um. **J.** MA-plot from microarray data published in Azami et al Development 2017 illustrating that, in this system, PrE marker genes are not significantly downregulated following loss of *Klf5.* Additionally, multiple probes for *Klf5* itself are not significantly differentially expressed.

**Supplementary Figure S4.**
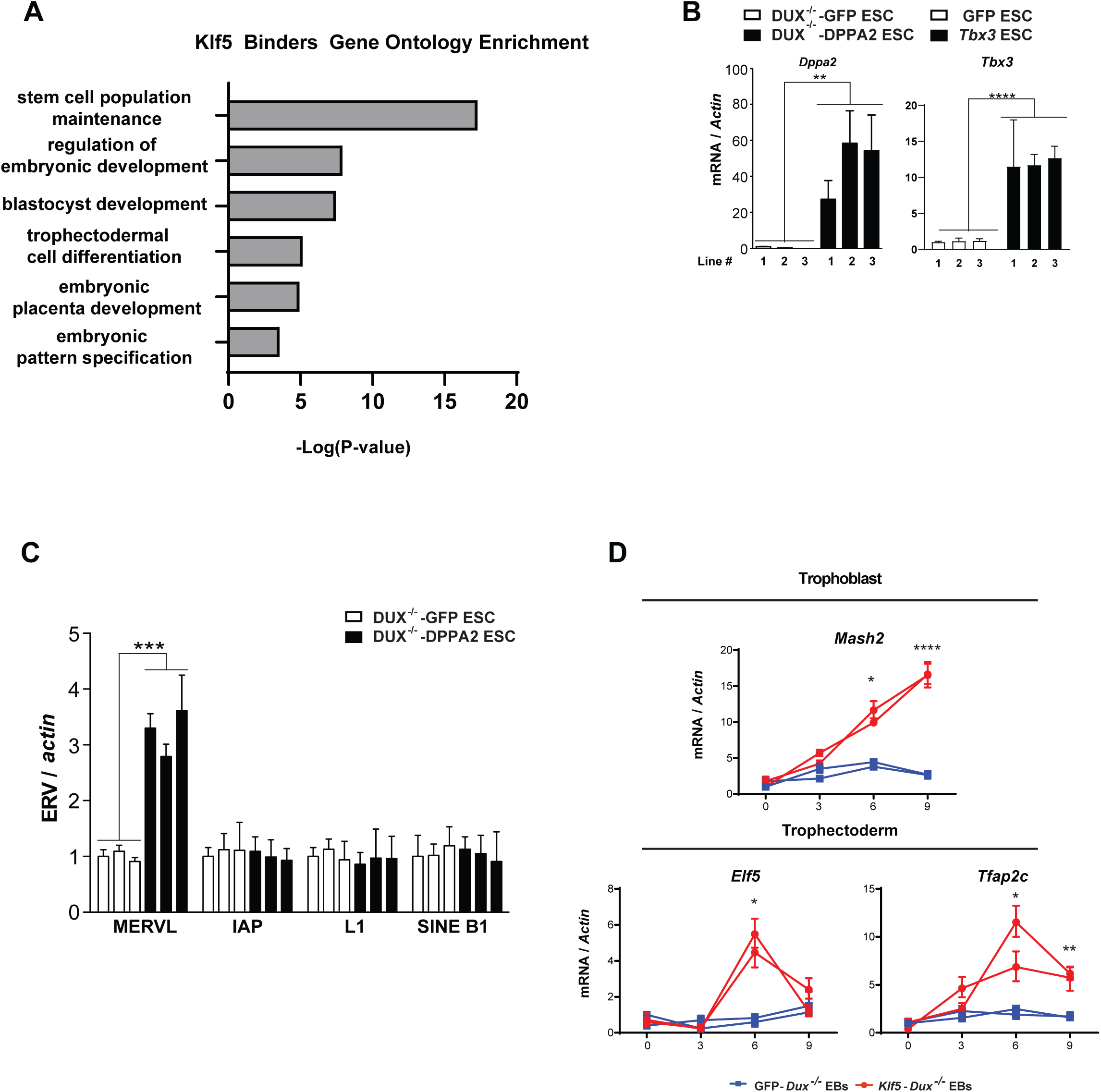
Klf5 induction is regulated by the 2C-specific transcription factor Dux. **A.** A gene ontology enrichment plot summarizing key enriched functional terms shared by *Klf5*-binding transcription factors. Specific processes in embryonic development and stem cell biology converge on the regulation of *Klf5*. **B.** Real time PCR analyses for *Dppa2* and *Klf5* in *Dppa2*-overexpressing and control *Dux*^-/-^ ESCs showing that *Klf5* is upregulated by *Dppa2* in ESCs, independently of *Dux*. Three independent *Dux*^-/-^ ESC lines were employed for this analysis. Error bars = s.d., *Dppa2*, \*\**P =* 0.0090, df = 4, t = 4.743; *Klf5*, \*\**P* = 0.0027, df = 4, t = 6.622. **C.** EBs derived from *Klf5-*overexpressing *Dux*^-/-^ ESCs induce markers of TE and trophoblasts. Real time PCR analyses confirmed the additional induction of TE lineage markers and trophoblast markers differentiating EBs derived from *Klf5*-overexpressing *Dux*^-/-^ ESCs, but not from control *Dux*^-/-^ ESCs. Two independent *Dux*^-/-^ ESC lines were employed for these analyses. Error bars = s.d. *Elf5* (day 6), * *P* = 0.0147, df = 2, t = 8.158; *Tfap2c* (day 6), * *P* = 0.0403, df = 2, t = 4.830; *Tfap2C* (day 9), ** *P* = 0.0014, df = 2, t = 26.55; *Mash2* (day 6), * *P* = 0.0132, df = 2, t = 8.602; *Mash2* (day 9), **** *P* < 0.0001, df = 2, t = 138.8. All P values were calculated on the basis of unpaired, two-tailed Student’s t-tests. * *P* < 0.05, ** *P* < 0.01, *** *P* < 0.001, **** *P* < 0.0001

## References and Notes

(IMPC), M.G.I. and the I.M.P.C. (2014). Obtaining and Loading Phenotype Annotations from the International Mouse Phenotyping Consortium (IMPC) Database. Database Release.

Alda-Catalinas, C., Bredikhin, D., Hernando-Herraez, I., Santos, F., Kubinyecz, O., Eckersley-Maslin, M.A., Stegle, O., and Reik, W. (2020). A Single-Cell Transcriptomics CRISPR-Activation Screen Identifies Epigenetic Regulators of the Zygotic Genome Activation Program. Cell Syst.

Azami, T., Waku, T., Matsumoto, K., Jeon, H., Muratani, M., Kawashima, A., Yanagisawa, J., Manabe, I., Nagai, R., Kunath, T., et al. (2017). Klf5 maintains the balance of primitive endoderm versus epiblast specification during mouse embryonic development by suppression of Fgf4. Dev.

Beddington, R.S.P., and Robertson, E.J. (1989). An assessment of the developmental potential of embryonic stem cells in the midgestation mouse embryo. Development.

Benner, C., Heinz, S., and Glass, C.K. (2017). HOMER - Software for motif discovery and next generation sequencing analysis. http://Homer.Ucsd.Edu/.

Bialkowska, A.B., Yang, V.W., and Mallipattu, S.K. (2017). Krüppel-like factors in mammalian stem cells and development. Dev.

Casser, E., Israel, S., Witten, A., Schulte, K., Schlatt, S., Nordhoff, V., and Boiani, M. (2017). Totipotency segregates between the sister blastomeres of two-cell stage mouse embryos. Sci. Rep. 7.

Chen, Z., and Zhang, Y. (2019). Loss of DUX causes minor defects in zygotic genome activation and is compatible with mouse development. Nat. Genet.

Choi, Y.J., Lin, C.-P., Risso, D., Chen, S., Kim, T.A., Tan, M.H., Li, J.B., Wu, Y., Chen, C., Xuan, Z., et al. (2017). Deficiency of microRNA miR-34a expands cell fate potential in pluripotent stem cells. Science (80-.). 355, eaag1927.

Dai, Q., Shen, Y., Wang, Y., Wang, X., Francisco, J.C., Luo, Z., and Lin, C. (2017). Striking a balance: regulation of transposable elements by Zfp281 and Mll2 in mouse embryonic stem cells. Nucleic Acids Res.

Dan, J., Li, M., Yang, J., Li, J., Okuka, M., Ye, X., and Liu, L. (2013). Roles for Tbx3 in regulation of two-cell state and telomere elongation in mouse ES cells. Sci. Rep. 3, 3492.

Deng, Q., Ramsköld, D., Reinius, B., Sandberg, R., Ramskold, D., Reinius, B., and Sandberg, R. (2014). Single-Cell RNA-Seq Reveals Dynamic, Random Monoallelic Gene Expression in Mammalian Cells. Science (80-.). 343, 193–196.

Eckersley-Maslin, M., Alda-Catalinas, C., Blotenburg, M., Kreibich, E., Krueger, C., and Reik, W. (2019). Dppa2 and Dppa4 directly regulate the Dux-driven zygotic transcriptional program. Genes Dev.

Franke, V., Ganesh, S., Karlic, R., Malik, R., Pasulka, J., Horvat, F., Kuzman, M., Fulka, H., Cernohorska, M., Urbanova, J., et al. (2017). Long terminal repeats power evolution of genes and gene expression programs in mammalian oocytes and zygotes. Genome Res. 27, 1384–1394.

Fujimori, T., Kurotaki, Y., Miyazaki, J.I., and Nabeshima, Y.I. (2003). Analysis of cell lineage in two- and four-cell mouse embryos. Development.

Hendrickson, P.G., Doráis, J.A., Grow, E.J., Whiddon, J.L., Lim, J.-W.W., Wike, C.L., Weaver, B.D., Pflueger, C., Emery, B.R., Wilcox, A.L., et al. (2017). Conserved roles of mouse DUX and human DUX4 in activating cleavage-stage genes and MERVL/HERVL retrotransposons. Nat. Genet. 49, 925–934.

Hirate, Y., Hirahara, S., Inoue, K.I., Suzuki, A., Alarcon, V.B., Akimoto, K., Hirai, T., Hara, T., Adachi, M., Chida, K., et al. (2013). Polarity-dependent distribution of angiomotin localizes hippo signaling in preimplantation embryos. Curr. Biol. 23, 1181–1194.

Hu, Z., Tan, D.E.K., Chia, G., Tan, H., Leong, H.F., Chen, B.J., Lau, M.S., Tan, K.Y.S., Bi, X., Yang, D., et al. (2020). Maternal factor NELFA drives a 2C-like state in mouse embryonic stem cells. Nat. Cell Biol.

Hubley, R., Finn, R.D., Clements, J., Eddy, S.R., Jones, T.A., Bao, W., Smit, A.F.A., and Wheeler, T.J. (2016). The Dfam database of repetitive DNA families. Nucleic Acids Res.

Iaco, A. De, Planet, E., Coluccio, A., Verp, S., Duc, J., Trono, D., De Iaco, A., Planet, E., Coluccio, A., Verp, S., et al. (2017). DUX-family transcription factors regulate zygotic genome activation in placental mammals. Nat. Genet. 49, 941–945.

Ishiuchi, T., Enriquez-gasca, R., Mizutani, E., Boškoviä, A., Ziegler-birling, C., Rodriguez-terrones, D., Wakayama, T., Vaquerizas, J.M., Torres-Padilla, M.E., Bošković, A., et al. (2015a). Articles Early embryonic-like cells are induced by downregulating replication-dependent chromatin assembly. Nat. Publ. Gr. 22, 662–671.

Ishiuchi, T., Enriquez-Gasca, R., Mizutani, E., Boškoviä, A., Ziegler-Birling, C., Rodriguez-Terrones, D., Wakayama, T., Vaquerizas, J.M., and Torres-Padilla, M.E. (2015b). Early embryonic-like cells are induced by downregulating replication-dependent chromatin assembly. Nat. Struct. Mol. Biol.

Katz, J.P., Perreault, N., Goldstein, B.G., Lee, C.S., Labosky, P.A., Yang, V.W., and Kaestner, K.H. (2002). The zinc-finger transcription factor Klf4 is required for terminal differentiation of goblet cells in the colon. Development.

Kelly, S.J. (1977). Studies of the developmental potential of 4- and 8- cell stage mouse blastomeres. J. Exp. Zool.

Korotkevich, E., Niwayama, R., Courtois, A., Friese, S., Berger, N., Buchholz, F., and Hiiragi, T. (2017). The Apical Domain Is Required and Sufficient for the First Lineage Segregation in the Mouse Embryo. Dev. Cell 40, 235–247.e7.

Lin, S.-C.J., Wani, M.A., Whitsett, J.A., and Wells, J.M. (2010). Klf5 regulates lineage formation in the pre-implantation mouse embryo. Development 137, 3953–3963.

M., E., D., M., H., N., Y., H., Y., Y., S., H., J., M., Y.-i., K., T., H., M., M., et al. (2008). Krüppel-like factor 5 Is Essential for Blastocyst Development and the Normal Self-Renewal of Mouse ESCs. Cell Stem Cell 3, 555–567.

Macfarlan, T.S., Gifford, W.D., Agarwal, S., Driscoll, S., Lettieri, K., Wang, J., Andrews, S.E., Franco, L., Rosenfeld, M.G., Ren, B., et al. (2011). Endogenous retroviruses and neighboring genes are coordinately repressed by LSD1/KDM1A. Genes Dev. 25, 594–607.

Macfarlan, T.S., Gifford, W.D., Driscoll, S., Lettieri, K., Rowe, H.M., Bonanomi, D., Firth, A., Singer, O., Trono, D., and Pfaff, S.L. (2012). Embryonic stem cell potency fluctuates with endogenous retrovirus activity. Nature.

McCarthy, E.M., and McDonald, J.F. (2004). Long terminal repeat retrotransposons of Mus musculus. Genome Biol.

Mei, S., Qin, Q., Wu, Q., Sun, H., Zheng, R., Zang, C., Zhu, M., Wu, J., Shi, X., Taing, L., et al. (2017). Cistrome Data Browser: A data portal for ChIP-Seq and chromatin accessibility data in human and mouse. Nucleic Acids Res.

Nakamura, T., Nakagawa, M., Ichisaka, T., Shiota, A., and Yamanaka, S. (2011). Essential Roles of ECAT15-2/Dppa2 in Functional Lung Development. Mol. Cell. Biol.

Pfeffer, P.L. (2018). Building principles for constructing a mammalian blastocyst embryo. Biology (Basel).

Presnell, J.S., Schnitzler, C.E., and Browne, W.E. (2015). KLF/SP transcription factor family evolution: Expansion, diversification, and innovation in eukaryotes. Genome Biol. Evol.

Schoorlemmer, J., Pérez-Palacios, R., Climent, M., Guallar, D., and Muniesa, P. (2014). Regulation of Mouse Retroelement MuERV-L/MERVL Expression by REX1 and Epigenetic Control of Stem Cell Potency. Front. Oncol. 4, 14.

Stopka, T., and Skoultchi, A.I. (2003). The ISWI ATPase Snf2h is required for early mouse development. Proc. Natl. Acad. Sci. U. S. A.

Strumpf, D. (2005). Cdx2 is required for correct cell fate specification and differentiation of trophectoderm in the mouse blastocyst. Development 132, 2093–2102.

Tabansky, I., Lenarcic, A., Draft, R.W., Loulier, K., Keskin, D.B., Rosains, J., Rivera-Feliciano, J., Lichtman, J.W., Livet, J., Stern, J.N.H., et al. (2013). Developmental bias in cleavage-stage mouse blastomeres. Curr. Biol.

Tagliaferri, D., Mazzone, P., Noviello, T.M.R., Addeo, M., Angrisano, T., Del Vecchio, L., Visconte, F., Ruggieri, V., Russi, S., Caivano, A., et al. (2020). Retinoic Acid Induces Embryonic Stem Cells (ESCs) Transition to 2 Cell-Like State Through a Coordinated Expression of Dux and Duxbl1. Front. Cell Dev. Biol.

Wigger, M., Kisielewska, K., Filimonow, K., Plusa, B., Maleszewski, M., and Suwińska, A. (2017). Plasticity of the inner cell mass in mouse blastocyst is restricted by the activity of FGF/MAPK pathway. Sci. Rep.

Wu, G., Lei, L., and Schöler, H.R. (2017). Totipotency in the mouse. J. Mol. Med.

Yamane, M., Ohtsuka, S., Matsuura, K., Nakamura, A., and Niwa, H. (2018). Overlapping functions of krüppel-like factor family members: Targeting multiple transcription factors to maintain the naïve pluripotency of mouse embryonic stem cells. Dev.

Yan, Y.L., Zhang, C., Hao, J., Wang, X.L., Ming, J., Mi, L., Na, J., Hu, X., and Wang, Y. (2019). DPPA2/4 and SUMO E3 ligase PIAS4 opposingly regulate zygotic transcriptional program. PLoS Biol.

Yang, J., Ryan, D.J., Wang, W., Tsang, J.C.H., Lan, G., Masaki, H., Gao, X., Antunes, L., Yu, Y., Zhu, Z., et al. (2017a). Establishment of mouse expanded potential stem cells. Nature 550, 393–397.

Yang, Y., Liu, B., Xu, J., Wang, J., Wu, J., Shi, C., Xu, Y., Dong, J., Wang, C., Lai, W., et al. (2017b). Derivation of Pluripotent Stem Cells with In Vivo Embryonic and Extraembryonic Potency. Cell 169, 243–257.e25.

Yeo, J.C., Jiang, J., Tan, Z.Y., Yim, G.R., Ng, J.H., Göke, J., Kraus, P., Liang, H., Gonzales, K.A.U., Chong, H.C., et al. (2014). Klf2 is an essential factor that sustains ground state pluripotency. Cell Stem Cell.

Zhao, T., Fu, Y., Zhu, J., Liu, Y., Zhang, Q., Yi, Z., Chen, S., Jiao, Z., Xu, X., Xu, J., et al. (2018). Single-Cell RNA-Seq Reveals Dynamic Early Embryonic-like Programs during Chemical Reprogramming. Cell Stem Cell 23, 31–45.e7.

## Supplementary references

Anders, S., and Huber, W. (2010). Differential expression analysis for sequence count data. Genome Biol.

Benner, C., Heinz, S., and Glass, C.K. (2017). HOMER - Software for motif discovery and next generation sequencing analysis.

Van den Berge, K., Roux de Bézieux, H., Street, K., Saelens, W., Cannoodt, R., Saeys, Y., Dudoit, S., and Clement, L. (2020). Trajectory-based differential expression analysis for single-cell sequencing data. Nat. Commun. 11, 1201.

Bray, N.L., Pimentel, H., Melsted, P., and Pachter, L. (2016). Near-optimal probabilistic RNA-seq quantification. Nat. Biotechnol. 34, 525–527.

Bryja, V., Bonilla, S., and Arenas, E. (2006). Derivation of mouse embryonic stem cells. Nat. Protoc.

Choi, Y.J., Lin, C.P., Ho, J.J., He, X., Okada, N., Bu, P., Zhong, Y., Kim, S.Y., Bennett, M.J., Chen, C., et al. (2011). MiR-34 miRNAs provide a barrier for somatic cell reprogramming. Nat. Cell Biol.

Choi, Y.J., Lin, C.P., Risso, D., Chen, S., Kim, T.A., Tan, M.H., Li, J.B., Wu, Y., Chen, C., Xuan, Z., et al. (2017). Deficiency of microRNA miR-34a expands cell fate potential in pluripotent stem cells. Science (80-.). 355.

Dobin, A., Davis, C.A., Schlesinger, F., Drenkow, J., Zaleski, C., Jha, S., Batut, P., Chaisson, M., and Gingeras, T.R. (2013). STAR: Ultrafast universal RNA-seq aligner. Bioinformatics 29, 15–21.

Harrow, J., Frankish, A., Gonzalez, J.M., Tapanari, E., Diekhans, M., Kokocinski, F., Aken, B.L., Barrell, D., Zadissa, A., Searle, S., et al. (2012). GENCODE: The reference human genome annotation for the ENCODE project. Genome Res. 22, 1760–1774.

Hendrickson, P.G., Doráis, J.A., Grow, E.J., Whiddon, J.L., Lim, J.W., Wike, C.L., Weaver, B.D., Pflueger, C., Emery, B.R., Wilcox, A.L., et al. (2017). Conserved roles of mouse DUX and human DUX4 in activating cleavage-stage genes and MERVL/HERVL retrotransposons. Nat. Genet. 49, 925–934.

Liao, Y., Smyth, G.K., and Shi, W. (2014). FeatureCounts: An efficient general purpose program for assigning sequence reads to genomic features. Bioinformatics 30, 923–930.

Mathelier, A., Fornes, O., Arenillas, D.J., Chen, C.Y., Denay, G., Lee, J., Shi, W., Shyr, C., Tan, G., Worsley-Hunt, R., et al. (2016). JASPAR 2016: A major expansion and update of the open-access database of transcription factor binding profiles. Nucleic Acids Res.

Modzelewski, A.J., Chen, S., Willis, B.J., Lloyd, K.C.K., Wood, J.A., and He, L. (2018). Efficient mouse genome engineering by CRISPR-EZ technology. Nat. Protoc.

Ritchie, M.E., Phipson, B., Wu, D., Hu, Y., Law, C.W., Shi, W., and Smyth, G.K. (2015). limma powers differential expression analyses for RNA-sequencing and microarray studies. Nucleic Acids Res. 43, e47–e47.

Street, K., Risso, D., Fletcher, R.B., Das, D., Ngai, J., Yosef, N., Purdom, E., and Dudoit, S. (2018). Slingshot: Cell lineage and pseudotime inference for single-cell transcriptomics. BMC Genomics 19.

Wang, W.C.H., and Shashikant, C.S. (2007). Evidence for positive and negative regulation of the mouse Cdx2 gene. J. Exp. Zool. Part B Mol. Dev. Evol.

Zhang, Y., Liu, T., Meyer, C.A., Eeckhoute, J., Johnson, D.S., Bernstein, B.E., Nussbaum, C., Myers, R.M., Brown, M., Li, W., et al. (2008). Model-based analysis of ChIP-Seq (MACS). Genome Biol. 9, R137.

